# Small leucine-rich proteoglycans inhibit CNS regeneration by modifying the structural and mechanical properties of the lesion environment

**DOI:** 10.1101/2022.11.21.517128

**Authors:** Julia Kolb, Nora John, Kyoohyun Kim, Conrad Möckel, Gonzalo Rosso, Stephanie Möllmert, Veronika Kurbel, Asha Parmar, Gargi Sharma, Timon Beck, Paul Müller, Raimund Schlüßler, Renato Frischknecht, Anja Wehner, Nicole Krombholz, Barbara Steigenberger, Ingmar Blümcke, Kanwarpal Singh, Jochen Guck, Katja Kobow, Daniel Wehner

**Affiliations:** Max Planck Institute for the Science of Light, 91058 Erlangen, Germany; Max-Planck-Zentrum für Physik und Medizin, 91058 Erlangen, Germany; Department of Biology, Animal Physiology, Friedrich-Alexander-University Erlangen-Nürnberg, 91058 Erlangen, Germany; Department of Neuropathology, Universitätsklinikum Erlangen, Friedrich-Alexander-University Erlangen-Nürnberg, 91054 Erlangen, Germany; Department of Physics, Friedrich-Alexander-University Erlangen-Nürnberg, 91058 Erlangen, Germany; Department of Medicine 1, Universitätsklinikum Erlangen, Friedrich-Alexander-University Erlangen-Nürnberg, 91054 Erlangen, Germany; Biotechnology Center, Center for Molecular and Cellular Bioengineering, Technische Universität Dresden, 01307 Dresden, Germany; Mass Spectrometry Core Facility, Max Planck Institute of Biochemistry, 82152 Martinsried, Germany

**Keywords:** CNS injury, axon regeneration, ECM, small leucine-rich proteoglycans, Brillouin microscopy, cross-polarized optical coherence tomography, optical diffraction tomography, tissue mechanics, scar

## Abstract

Extracellular matrix (ECM) deposition after central nervous system (CNS) injury leads to inhibitory scarring in mammals, whereas it facilitates axon regeneration in the zebrafish. However, the molecular basis of these different fates is not understood. Here, we identify small leucine-rich proteoglycans (SLRPs) as a causal factor in regeneration failure. We demonstrate that the SLRPs Chondroadherin, Fibromodulin, Lumican, and Prolargin are enriched in human, but not zebrafish, CNS lesions. Targeting SLRPs to the zebrafish injury ECM inhibits axon regeneration and functional recovery. Mechanistically, we find that SLRPs confer structural and mechanical properties to the lesion environment that are adverse to axon growth. Our study reveals SLRPs as previously unknown inhibitory ECM factors in the human CNS that impair axon regeneration by modifying tissue mechanics and structure.

**ONE SENTENCE SUMMARY:** Composition, structural organization, and mechanical properties of the injury ECM direct central nervous system regeneration.

## INTRODUCTION

The ability to regenerate long-distance axonal connections after central nervous system (CNS) injury differs significantly among vertebrates. Why certain species, such as zebrafish, possess a high regenerative capacity, but not others, is poorly understood (*1*). Fibrous scar formation is considered to be a major factor in limiting axon regeneration in the adult mammalian CNS (*2*). Several scar components, such as myelin-associated factors, basal lamina components, and high molecular weight chondroitin sulfate proteoglycans (CSPGs), have been identified as inhibitors of axonal regrowth through mechanisms including growth cone collapse, repellence or entrapment, and prevention of inflammation resolution (*3–5*). However, removing these extracellular matrix (ECM) factors results in only modest regeneration, suggesting that critical molecules and mechanisms contributing to regeneration failure remain to be discovered (*6–8*). The CNS scar may inhibit axon growth not only through its biochemical composition, but also by changes in local mechanical properties of the microenvironment (*9–11*). However, *in vivo* evidence for a causal relationship between scar tissue mechanics and regenerative success is lacking. Moreover, molecular factors that influence the mechanical properties of CNS scars have not been identified.

Unlike mammals, zebrafish establish an axon growth-conducive ECM after CNS injury, leading to recovery of swimming function both at larval and adult stages (*12–16*). It remains obscure as to why injury-associated ECM deposits inhibit axon regeneration in the mammalian CNS, but not in zebrafish. Here, we identify small leucine-rich proteoglycans (SLRPs) as ECM factors which drive CNS healing toward inhibitory scarring. We demonstrate that Chondroadherin, Fibromodulin, Lumican, and Prolargin are enriched in human but not zebrafish CNS lesions. Increasing the abundance of SLRPs in the zebrafish injury ECM inhibits axon regeneration and functional recovery. Mechanistically, we find that SLRPs confer structural and mechanical properties to the lesion environment that render it adverse to axon growth. This identifies SLRPs as previously unknown inhibitory ECM factors in the human CNS that impair axon regeneration by altering tissue mechanics and structure. Targeting SLRPs therefore presents itself as a potential therapeutic strategy to promote axon growth across CNS lesions.

## RESULTS

### Matrisome dynamics of zebrafish spinal cord regeneration

In order to identify factors that confer axon growth-limiting properties to CNS scars by altering tissue mechanics, we first set out to map the changes in ECM composition in a regeneration context. We applied label-free mass spectrometry (MS)-based quantitative proteomics to a larval zebrafish spinal cord injury (SCI) model, which allows axon regeneration and functional recovery within two days post-lesion (dpl) (Fig. 1A,B) (*13, 17*). Proteomic profiling of the lesion site at 1 dpl and 2 dpl as well as corresponding age-matched unlesioned control tissue identified 6,062 unique proteins (Fig. S1A,B). Differential abundance analysis (FDR < 0.1, |FC| ≥ 1.3, s_0_ = 0.1) revealed 877 proteins whose abundance was altered in 1 dpl compared to unlesioned control samples (556 up- and 321 down-regulated; Fig. S1C). The abundance of 570 proteins differed between 2 dpl and unlesioned control samples (388 up- and 182 down-regulated; Fig. S1D). Reactome analysis of enriched proteins resulted in several ECM-associated terms being overrepresented at 1 dpl and 2 dpl (Fig. S1E). We thus examined the matrisome, which can be subdivided into core matrisome (glycoproteins, collagens, proteoglycans), matrisome-associated (ECM affiliated proteins, ECM regulators, secreted factors), and putative matrisome proteins (Fig. 1C,D) (*18*). Among those matrisome proteins which exhibited a high abundance at 1 dpl were several proteins that have previously been reported to show increased expression after SCI in zebrafish, including Aspn, Cthrc1a, Col5a1, Col6a2, Col12a1a, Col12a1b, Fn1a, Fn1b, Tnc, and Thbs2b (*12–14, 19*). In addition, we identified immune system-related factors, such as cathepsins, serpins, galectins, and interleukins, which is consistent with the critical role of injury-activated macrophages at this stage of regeneration (*20, 21*). At 2 dpl, the number of matrisome proteins exhibiting an altered abundance decreased by 28%, compared to 1 dpl. Comparing differentially regulated proteins across time points showed that 30 such matrisome proteins were common to 1 dpl and 2 dpl (asterisks in Fig. 1C,D), including proteins previously implicated in axon growth promotion or guidance: Col12a1a, Col12a1b, Fn1a, Fn1b, Tnc, Cthrc1a, and Thbs1a (*12, 13, 19, 22–28*). 34 proteins with altered abundance were unique to 1 dpl, and 16 proteins to 2 dpl. Among the uniquely differentially regulated matrisome proteins at 1 dpl, we identified nine immune system-related and coagulation factors (Ctsa, Ctsla, Cts, F2, Il16, Lgals9l3, Plg, Serpind2fb, Serpine2), consistent with the initial blood clotting reaction and transient proinflammatory phase after SCI in zebrafish (*20, 29*). To further validate our MS results, we examined whether changes in mRNA levels accompanied the alterations in protein abundance. We performed *in situ* hybridization (ISH) to analyze the expression of genes coding for 30 differentially regulated matrisome proteins which exhibited high abundance at 1 dpl. Transcripts of all analyzed genes were locally upregulated in the lesion site compared to both adjacent unlesioned trunk tissue and unlesioned age-matched controls (Fig. S1F; data not shown). This reinforces the findings of our proteomics profiling. Collectively, these data identify the dynamics of the matrisome landscape after SCI in a vertebrate species which exhibits a high regenerative capacity for the CNS.

**Figure 1.**
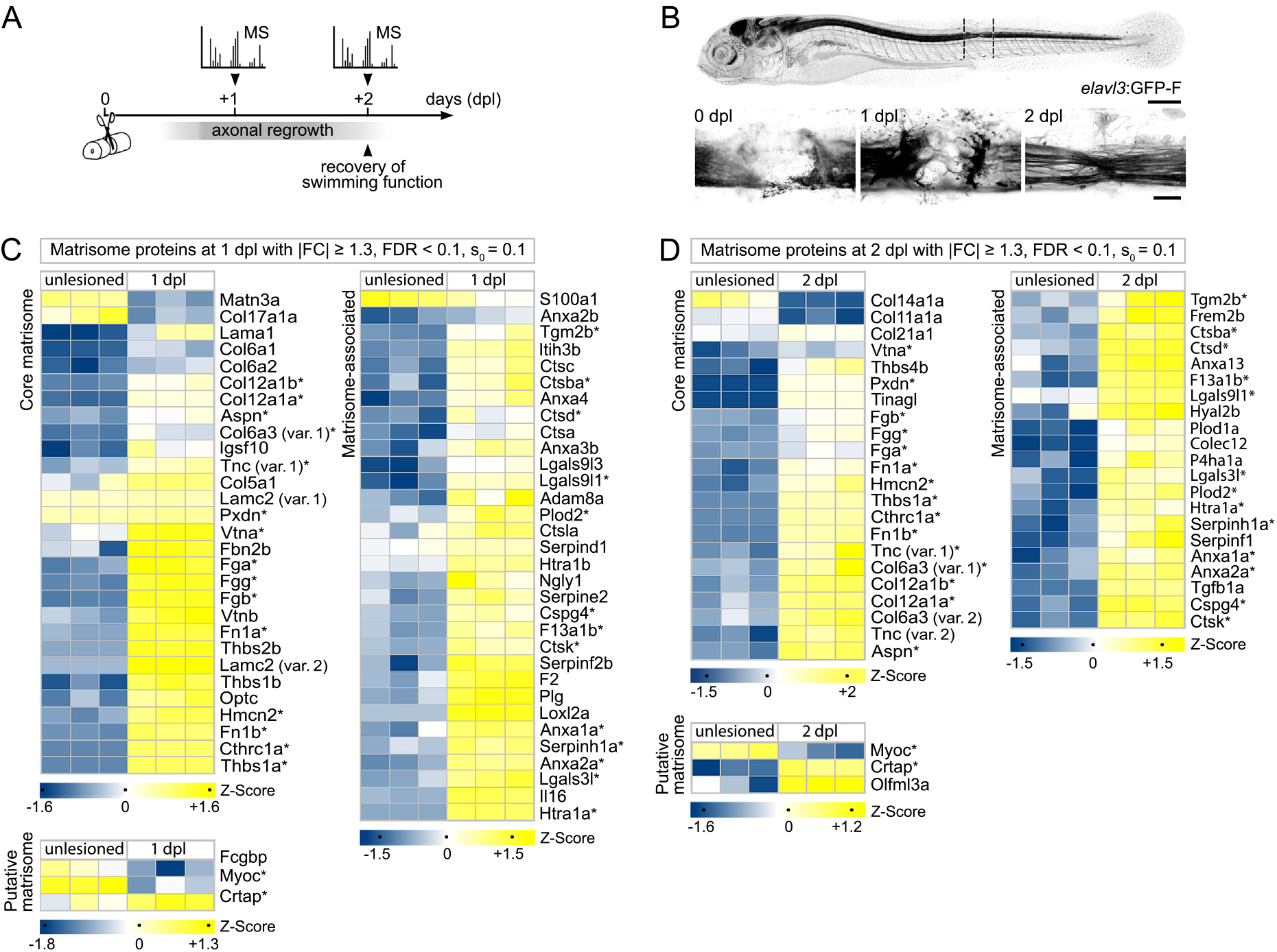
Mass spectrometry-based quantitative proteomics reveals changes in ECM composition during zebrafish spinal cord regeneration. **A)** Timeline of axonal regrowth and functional recovery after SCI in larval zebrafish. Timepoints of tissue collection for mass spectrometry (MS) analysis are indicated. **B)** Time course of axonal regrowth after spinal cord transection in *elavl3*:GFP-F transgenic zebrafish. Shown is the same animal at different timepoints after SCI. Dashed lines indicate the dissected trunk region for MS analysis. Images shown are maximum intensity projections of the spinal lesion site (lateral view; rostral is left). Scale bars: 250 μm (top) and 25 μm (bottom). **C-D)** Heatmaps of matrisome proteins exhibiting differential abundance between lesioned (1 dpl, C; 2 dpl, D) and unlesioned age-matched groups. Each column represents one biological replicate and each row one protein. Asterisks indicate matrisome proteins that are common to both timepoints. **A-D)** dpl, days post-lesion; FC, fold change; FDR, false discovery rate; var, variant.

### Distinct matrisome signatures after SCI in rat and zebrafish

To identify interspecies differences in the ECM composition that could account for the ability of severed axons to regrow after CNS injury, we compared the zebrafish proteomics dataset with that of Sprague-Dawley rats at seven days post-contusion SCI (*30*). We first screened for matrisome proteins that were enriched in the zebrafish lesion site (FDR < 0.1, FC ≥ 1.3) but down-regulated or not significantly regulated after SCI in rat (FDR < 0.1, FC ≤ −1.3 | n.s.). This identified four proteins (Fig. S2A). By contrast, 61 matrisome proteins were down-regulated or not significantly regulated after SCI in zebrafish (FDR < 0.1, FC ≤ −1.3 | n.s.) but enriched in the rat lesion site (FDR < 0.1, FC ≥ 1.3) (Fig. 2A). This suggests that the regeneration-permissive properties of the zebrafish injury ECM can be attributed to the absence of axon growth-limiting components rather than the presence of species-specific growth-promoting factors. Indeed, among the differentially regulated matrisome proteins that showed low abundance in the zebrafish spinal lesion site were 16 components of the neurite growth-inhibitory basal lamina (*4, 31, 32*). These include type IV collagens, laminins, nidogens, heparan sulfate proteoglycans, Fbln1, and Sparc. Quantitative RT-PCR (qRT-PCR) confirmed that the expression of these basal lamina components was not upregulated in the zebrafish spinal lesion site at 1 dpl (Fig. S2B). Similarly, anti-Col IV immunoreactivity was also not locally increased at 1 dpl (Fig. S2C). Thus, basal lamina networks are not a principal constituent of the zebrafish injury ECM. These data demonstrate the value of cross-species comparative approaches in identifying inhibitory components of the mammalian injury ECM. The comparative dataset further revealed seven members of the highly conserved small leucine-rich proteoglycan (SLRP) family to be enriched in the rat spinal lesion site but down-regulated or not significantly regulated after SCI in zebrafish, namely Chondroadherin (Chad), Lumican (Lum), Osteoglycin (Ogna), Decorin (Dcn), Fibromodulin (Fmoda, Fmodb), and Prolargin (Prelp) (Fig. 2A). Assessment of additional proteomics profiles showed comparable enrichment of these SLRPs in rats at seven days and eight weeks post-contusion SCI (Fig. S2D) (*30, 33, 34*). Interestingly, Asporin (Aspn) was the only SLRP family member which exhibited an increased abundance after SCI in both rat and zebrafish (Fig. S2D). To ascertain whether the low protein abundance of SLRPs in the zebrafish spinal lesion site is also reflected at the transcriptional level, we performed qRT-PCR. This revealed that with the exception of *aspn*, expression of all 21 SLRPs present in the zebrafish genome was not increased at 1 dpl as compared to unlesioned controls (Fig. 2B; Fig. S2E). Average fold changes of *aspn, chad, dcn, fmoda, fmodb, lum, ogna*, and *prelp*, as determined by qRT-PCR, correlated with the MS data (Pearson correlation, *R*^2^ = 0.9443), thus further validating our proteomics profile (Fig. S2F). Additionally, ISH showed upregulation of *aspn* transcripts in the lesion site at 1 dpl while *chad*, *dcn*, *fmoda*, *fmodb*, *lum, ogna*, and *prelp* expression was not locally increased (Fig. 2C; Fig. S2G). Altogether, these data reveal an opposing abundance of Chad, Lum, Ogn, Dcn, Fmod, and Prelp in the spinal lesion site of rat and zebrafish, respectively.

**Figure 2.**
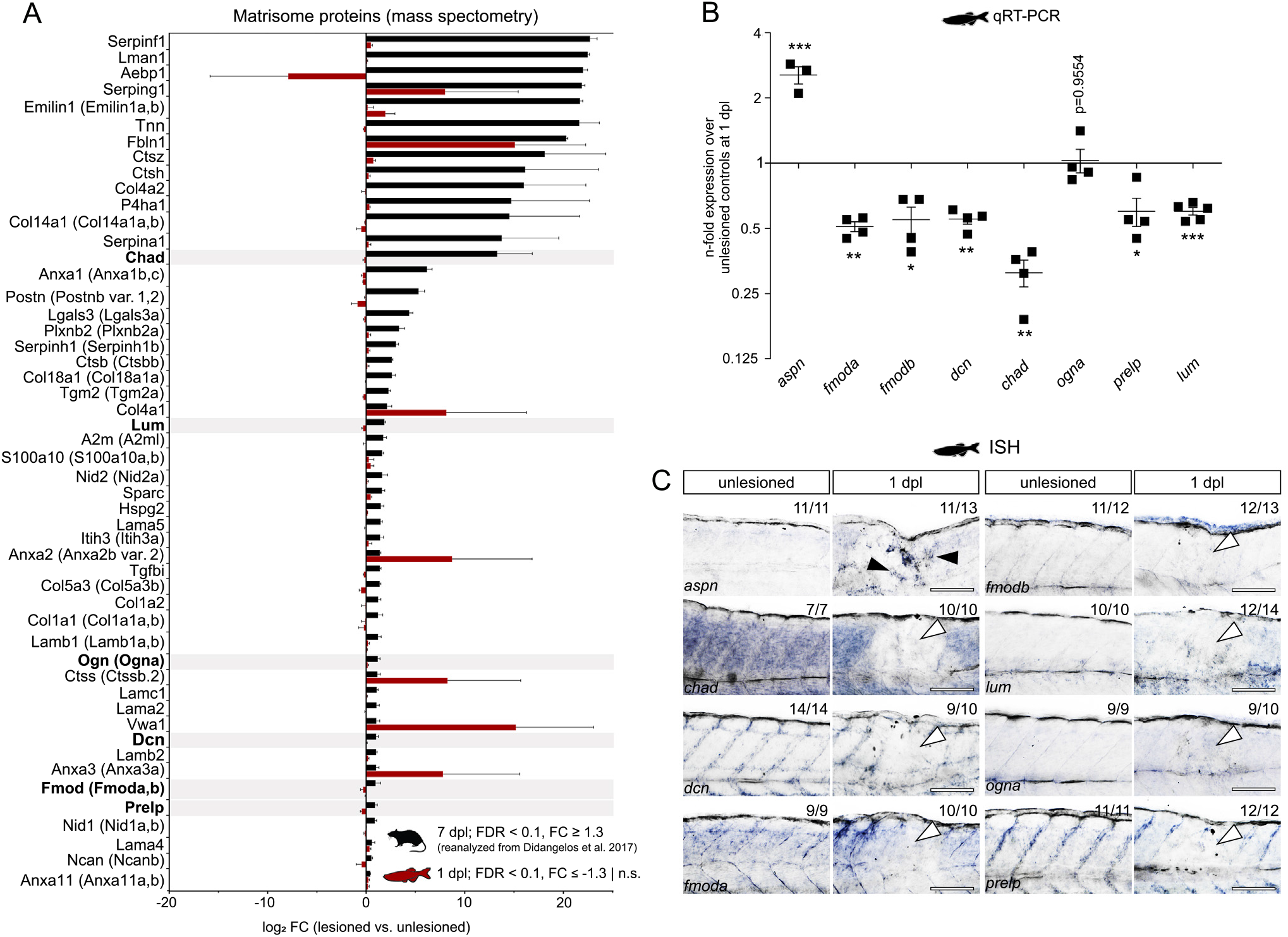
SLRPs are differentially enriched in CNS lesions of rat and zebrafish. **A)** Comparative proteomics analysis reveals differentially enriched matrisome proteins between rat (black) and zebrafish (red) after SCI, including the SLRPs Chad, Lum, Ogn (Ogna), Dcn, Fmod (Fmoda, Fmodb), and Prelp (labeled in bold). Proteins that exhibit a high abundance after SCI in rat but a low abundance in the zebrafish spinal lesion site are shown. Data are means ± SEM. **B)** Fold change expression of indicated genes in the zebrafish spinal lesion site at 1 dpl over unlesioned age-matched controls, as determined by qRT-PCR. Expression of *aspn* but not *fmoda*, *fmodb*, *dcn*, *chad*, *ogna, prelp*, and *lum* is upregulated at 1 dpl. Fold change values are presented in log scale. Each data point represents one biological replicate. Data are means ± SEM; \**P* < 0.05, \*\**P* < 0.01, \*\*\**P* < 0.001. **C)** Expression of *aspn* (black arrowheads) but not *chad, dcn, fmoda, fmodb, lum, ogna*, and *prelp* (white arrowheads) is upregulated in the zebrafish spinal lesion site at 1 dpl, as determined by *in situ* hybridization. The number of specimens displaying the phenotype and the total number of experimental specimens is given. Images shown are brightfield recordings of the lesion site or unlesioned trunk (lateral view; rostral is left). Scale bars: 100 μm. **A-C)** dpl, days post-lesion; FC, fold change; FDR, false discovery rate; n.s., not significant.

### SLRPs are abundant in human CNS lesions

We next sought to determine whether SLRPs are abundant in the injured human CNS. In surgically removed brain tissue samples from six patients with traumatic brain injury or brain surgery, we detected prominent anti-CHAD, anti-FMOD, anti-LUM, and anti-PRELP immunoreactivity in areas of scarring caused by contusion, local hemorrhage, or previous surgery (Fig. 3; Fig. S3A,B; Table S1). By contrast, negligible immunoreactivity was observed in a reference group of six human brain autopsy and biopsy controls without signs of fibrous scarring (Fig. S3C,D; Table S1). We next analyzed post-mortem spinal cord tissue from six patients with traumatic SCI. SCI occurred in the cervical region through compression or contusion, and patients survived between 9 and 111 days after injury (Table S2). We found a localized increase in immunoreactivity in the lesion epicenter, as compared to rostral or caudal control segments of the same patient, for anti-LUM in five out of six cases, for anti-PRELP and anti-FMOD in four out of six cases, and anti-CHAD in two out of six cases (Fig. S4; Table S3). The enrichment of SLRPs is therefore a feature of human CNS lesions.

**Figure 3.**
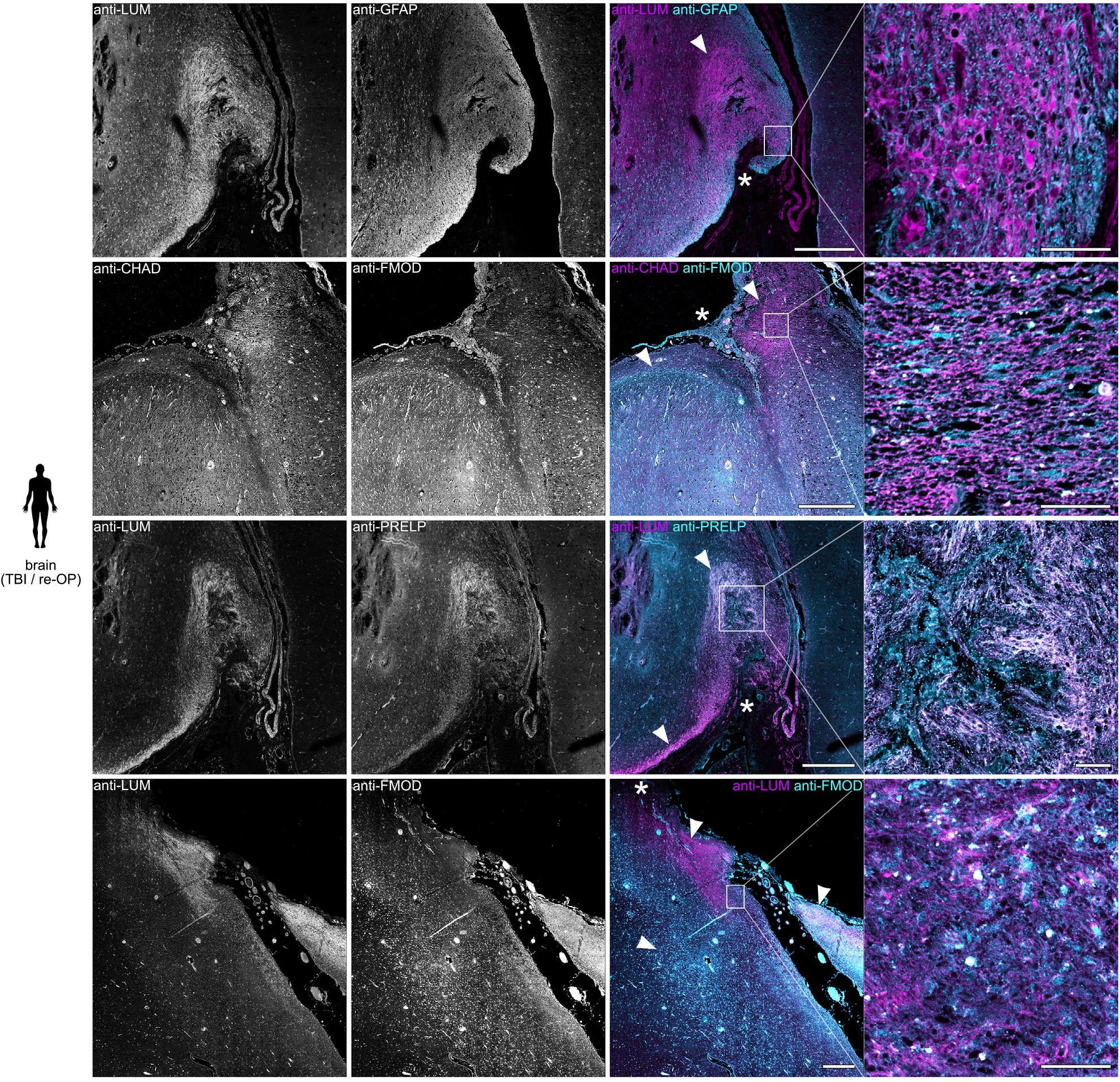
SLRPs are enriched in human brain lesions. Anti-CHAD, anti-FMOD, anti-LUM, and anti-PRELP immunoreactivity is increased (arrowheads) in areas of scarring caused by contusion, local hemorrhage (asterisk), or previous surgery in the human brain. Shown are coronal sections of brain tissue from patients with traumatic brain injury (TBI) or previous surgery (re-OP; bottom panel). Scale bars: 500 μm, 50 μm (insets).

### SLRPs are inhibitory to CNS axon regeneration

Since CHAD, FMOD, LUM, and PRELP proteins are highly abundant in the injury ECM of poorly regenerating humans but are absent in zebrafish with a high regenerative capacity for the CNS, we hypothesized that SLRPs inhibit regeneration. We therefore investigated the effect of increasing SLRP protein levels in the zebrafish injury ECM on axonal regrowth and functional recovery after SCI. We generated doxycycline (DOX)-inducible Tet-responder zebrafish lines to target *chad*, *fmoda*, *lum*, or *prelp* expression specifically to *pdgfrb*^+^ myoseptal and perivascular cells when used in combination with a *pdgfrb* promotor-driven Tet-activator line (henceforth also referred to as *pdgfrb*:SLRP). Additionally, we created a Tet-responder line for the selective induction of *aspn* to control for potential overexpression artifacts of SLRPs. *pdgfrb*^+^ fibroblast-like cells are rapidly recruited in response to SCI and constitute a major source of ECM in the lesion site (*12*). Moreover, qRT-PCR on GFP^+^ cells isolated from uninjured *pdgfrb*:GFP transgenic animals by FACS showed that *pdgfrb*^+^ cells express *aspn*, *chad*, *fmoda*, *lum*, and *prelp* under physiological conditions (Fig. S5A). This makes *pdgfrb*^+^ cells an ideal target for manipulating the ECM in the zebrafish spinal lesion site. Induction of *aspn-mCherry, chad-mCherry, fmoda-mCherry, lum-mCherry*, or *prelp-mCherry* fusions in *pdgfrb*:SLRP transgenics resulted in labeling of the myosepta and vasculature in unlesioned animals (Fig. S5B). At 1 dpl, mCherry fluorescence signal accumulated in the lesion site, indicating enrichment of the secreted proteins (Fig. 4A). *pdgfrb*:SLRP transgenics thereby enable the experimental increase of SLRP protein levels in the injury ECM after SCI. To assess whether axon regeneration is affected in *pdgfrb*:SLRP transgenics, we determined the thickness of the axonal bridge that reconnects the severed spinal cord ends in live *elavl3*:GFP-F transgenic animals at 2 dpl, a measure that correlates with the recovery of swimming function (Fig. 4B) (*20*). We found that the average axonal bridge thickness was reduced by 41-52% when Chad, Fmoda, Lum, or Prelp was targeted to the injury ECM, as compared to their controls (Fig. 4B’; Fig. S5C). Moreover, quantification of swimming distance at 2 dpl showed that *pdgfrb*:SLRP transgenics exhibited worse functional recovery than their controls (30-42% reduced swimming distance) (Fig. 4B”). Notably, targeting Aspn to the injury ECM had no effect on axon regeneration and recovery of swimming function, thus supporting the specificity of the observed phenotypes (Fig. 4B’,B”; Fig. S5C). Our data therefore identify Chad, Fmod, Lum, and Prelp as ECM factors that inhibit CNS axon regeneration *in vivo*.

**Figure 4.**
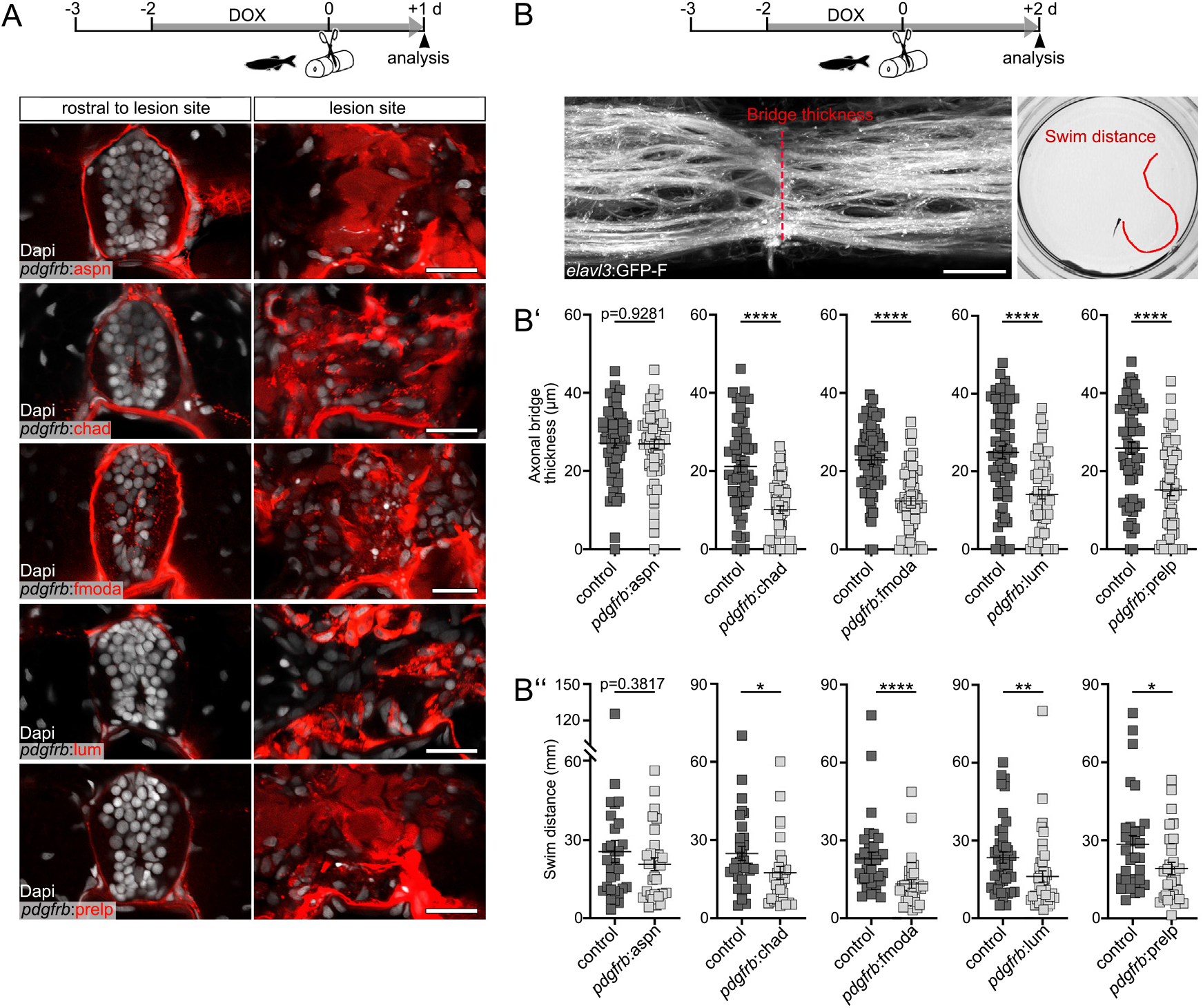
SLRPs are inhibitory to CNS axon regeneration. **A)** *pdgfrb*^+^ cell-specific induction of indicated *slrp*-*mCherry* fusions in *pdgfrb*:TetA;*TetRE*:SLRP-mCherry (short *pdgfrb*:SLRP) transgenic zebrafish leads to increased mCherry fluorescence (red) in the spinal lesion site at 1 dpl. Images shown are transversal views of the unlesioned trunk or lesion site (dorsal is up). **B)** *pdgfrb*^+^ cell-specific induction of the SLRPs *chad, fmoda, lum*, and *prelp* but not *aspn* in *pdgfrb*:SLRP transgenic zebrafish reduces the thickness of the axonal bridge (B’; analyzed in *elavl3*:GFP-F transgenics) and impairs recovery of swimming distance (B’’) at 2 dpl. Each data point represents one animal. Data are means ± SEM; \**P* < 0.05, \*\**P* < 0.01, \*\*\*\**P* < 0.0001. **A-B)** Scale bars: 20 μm. d, days; dpl, days post-lesion; DOX, doxycycline.

### SLRPs do not directly act on neurons to inhibit neurite extension

To elucidate the mechanism by which SLRPs inhibit neurite growth, we first examined a potential direct interaction with neurons or the axonal growth cone. We used a *Xla.Tubb* promoter-driven Tet-activator zebrafish line to target *chad*, *fmoda*, *lum*, or *prelp* expression specifically to neurons (referred to as *Xla.Tubb*:SLRP). In uninjured animals, the SLRP-mCherry fusion proteins were mainly confined to the spinal cord and showed negligible fluorescence signal in the lesion site at 1 dpl (Fig. S5D,E). Thus, cell type-specific manipulations allow us to distinguish between direct and indirect actions of SLRPs on axon growth *in vivo*. We found that targeting *chad*, *fmoda*, *lum*, or *prelp* to neurons did not impair axon regeneration, determined by measuring the axonal bridge thickness in live *elavl3:GFP-* F transgenic animals at 2 dpl (Fig. S5F). To corroborate these findings, we assessed neurite outgrowth of adult primary murine dorsal root ganglion neurons on SLRP protein-coated substrates. Although a mixture of high molecular weight CSPGs potently reduced the average neurite length by 66%, a combination of human CHAD, FMOD, LUM, and PRELP proteins had no effect (Fig. S5G). These data demonstrate that SLRPs inhibit axon growth indirectly.

### SLRPs do not prevent inflammation resolution

In order to test whether the presence of SLRPs in the injury ECM delays inflammation resolution, we analyzed the clearance of neutrophils and expression of the proinflammatory cytokine *il1b*, both of which processes must be tightly controlled for successful axon regeneration (*3, 20*). In larval zebrafish, the influx of Mpx^+^ neutrophils peaks as early as two hours following SCI, after which cell numbers rapidly decrease in the lesion site (*20*). We found that targeting Chad, Fmoda, Lum, or Prelp to the injury ECM in *pdgfrb*:SLRP transgenic animals did not lead to a higher number of Mpx^+^ neutrophils in the lesion site at 1 dpl (Fig. S6A). Consistent with this, transcript levels of neutrophil-derived *il1b* were not increased in the lesion site of *pdgfrb*:SLRP transgenic animals at 1 dpl (determined by qRT-PCR; Fig. S6B) (*20*). Altogether, this suggests that the inhibition of axon regeneration by SLRPs does not occur via the prevention of inflammation resolution.

### SLRPs do not alter the fibroblast response

We next sought to determine whether SLRPs inhibit axon regeneration by altering the fibroblast response in *pdgfrb*:SLRP transgenic animals. Using the TUNEL assay, we found that induction of *chad, fmoda, lum*, or *prelp* did not lead to increased cell death of *pdgfrb*^+^ cells (Fig. S7A,B). Furthermore, we did not detect significant changes in the area coverage of *pdgfrb*^+^ fibroblast-like cells in the lesion site at 1 dpl, indicating that their recruitment was largely unperturbed (Fig. S7C). Finally, we did not detect differences in the composition of the injury ECM between *pdgfrb*:SLRP transgenics and controls at 1 dpl (FDR < 0.1, |FC| ≥ 1.3), as revealed by MS-based quantitative proteomics (Fig. S8A-C). To further validate the MS results, we used ISH to evaluate the expression pattern and transcript levels of five genes coding for matrisome proteins that showed the most substantial evidence of regulation in each experimental condition. Consistent with the MS analysis, ISH signals in the lesion site were comparable between controls and *pdgfrb*:SLRP transgenics for all 25 genes analyzed (Fig. S8D). We thus conclude that targeting SLRPs to *pdgfrb*^+^ fibroblasts does not lead to major changes in the biochemical composition of the injury ECM.

### Fmod, Lum, and Prelp modify the structural properties of the lesion environment

SLRPs play instructive and structural roles in ECM organization and assembly to control the strength and biomechanical properties of tissues (*35*). We thus hypothesized that SLRPs alter the structural and mechanical properties of the injury ECM, thereby making it hostile to axon regeneration. In order to assess this *in vivo*, we first utilized cross-polarized optical coherence tomography (CP-OCT). CP-OCT reports on relative changes in the polarization of incident light and can provide additional contrast to the native tissue based on its structural differences, including ECM structure (*36, 37*). Consistent with the changes in tissue structure which occur after CNS injury, the co-polarization ratio (ratio of preserved polarization to total reflectivity) prominently increased in the zebrafish spinal lesion site at 1 dpl, as compared to adjacent uninjured trunk tissue (Fig. 5A,B). Targeting Chad to the injury ECM (analyzed in *pdgfrb*:SLRP transgenics) did not alter the co-polarization ratio in the lesion site at 1 dpl, as compared to controls (Fig. 5B). By contrast, an increase in the co-polarization ratio was observed when Fmoda, Lum, or Prelp were targeted to the injury ECM (control^Fmoda^: 0.709 ± 0.008, Fmoda: 0.750 ± 0.006; control^Lum^: 0.702 ± 0.010, Lum: 0.733 ± 0.005; control^Prelp^: 0.713 ± 0.006, Prelp: 0.738 ± 0.004; Fig. 5B). Hence, the presence of Fmod, Lum, and Prelp in the injury ECM coincides with alterations in the structural properties of the lesion environment and impaired axonal regrowth after SCI.

**Figure 5.**
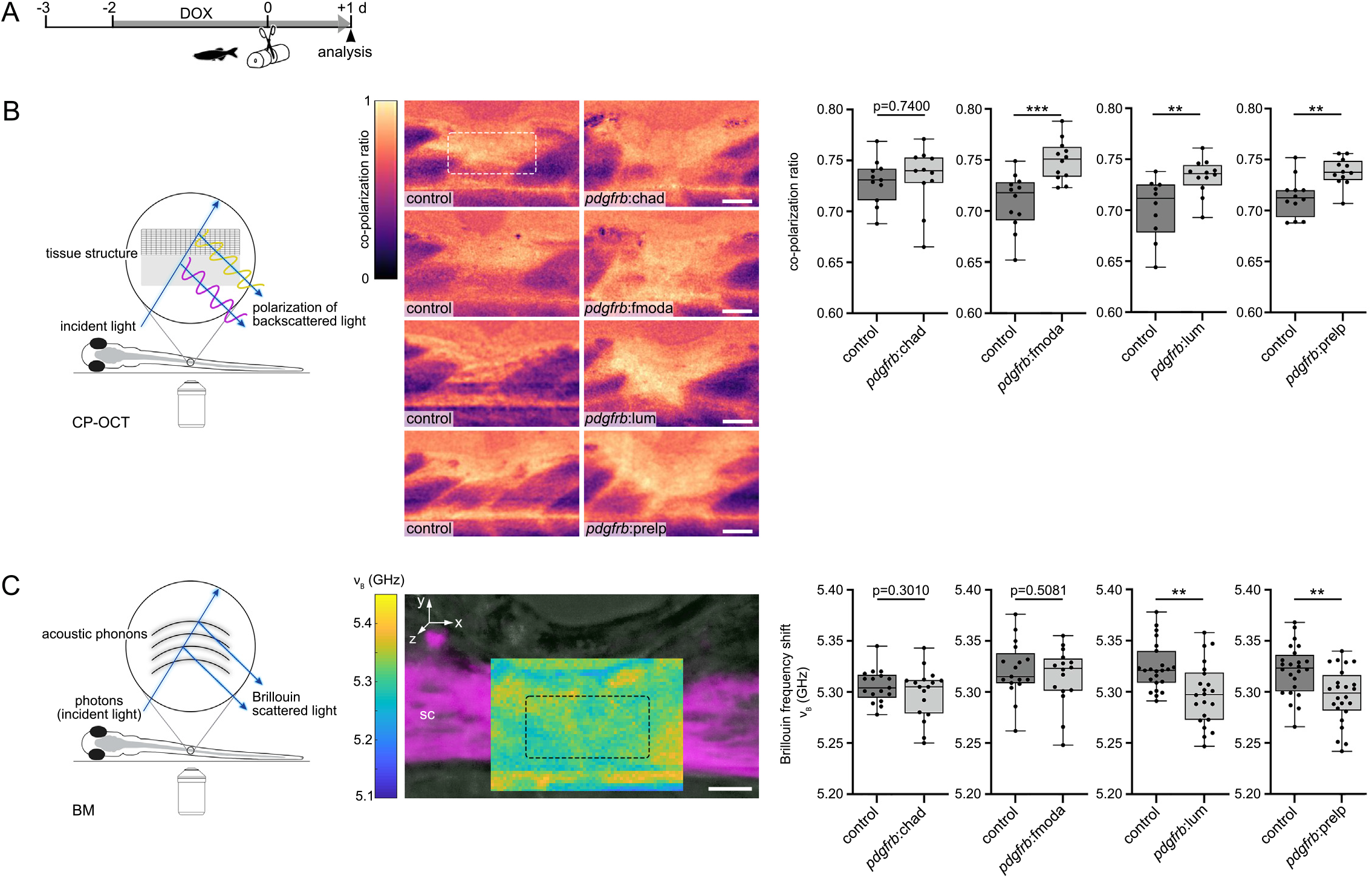
SLRPs modulate the structural and mechanical properties of the lesion environment. **A)** Timeline for experimental treatments shown in (B) and (C). **B)** Targeting Fmoda, Lum, or Prelp to the injury ECM in *pdgfrb*:SLRP transgenic zebrafish increases the co-polarization ratio (ratio of preserved polarization to total reflectivity) in the spinal lesion site, as determined by cross-polarized optical coherence tomography (CP-OCT) at 1 dpl. Images shown are average intensity projections of the lesion site (lateral view; rostral is left). **C)** Targeting Lum or Prelp to the injury ECM in *pdgfrb*:SLRP transgenic zebrafish decreases the mean Brillouin frequency shift (*ν*_B_) in the spinal lesion site, as determined by Brillouin microscopy (BM). Image shown is a sagittal optical section (overlay of brightfield intensity, confocal fluorescence, Brillouin frequency shift map) through the center of the lesion site of an *elavl3*:GFP-F transgenic zebrafish at 1 dpl (lateral view; rostral is left). **B-C)** The dashed rectangle indicates the region of quantification. Each data point represents one animal. Box plots show the median, first and third quartile. Whiskers indicate the minimum and maximum values. \*\**P* < 0.01, \*\*\**P* < 0.001. Scale bars: 50 μm (B), 25 μm (C). d, days; dpl, days post-lesion; DOX, doxycycline; sc, spinal cord.

### Lum and Prelp modify the mechanical properties of the lesion environment

We next explored whether the observed structural alterations relate to changes in the mechanical properties of the lesion environment. To test this *in vivo*, we utilized confocal Brillouin microscopy (BM), a non-invasive, label-free, and all-optical method for assessing viscoelastic properties of biological samples in three dimensions (*38–40*). BM measures an inelastic scattering process of incident light (photons) by density fluctuations (acoustic phonons) in the sample, called Brillouin scattering. The energy transfer between the photons and acoustic phonons occurring during the scattering process can be quantified as the Brillouin frequency shift (*ν*_B_), which depends on the material’s elastic properties, denoted by the longitudinal modulus (*M*’). *ν*_B_ is proportional to the square root of *M*’, which describes a material’s elastic deformability under a distinct type of mechanical loading (i.e., longitudinal compressibility). Notably, *ν*_B_ has been shown *in vivo* to be sensitive to changes in the mechanical properties of the ECM during both physiological and pathological processes (*41–43*). We acquired Brillouin images of the region in the spinal lesion site through which the regenerating axons preferentially grow at 1 dpl (Fig. 5A,C). Targeting Chad or Fmoda to the injury ECM (analyzed in *pdgfrb*:SLRP transgenics) did not alter *ν*_B_ of the lesion site (Fig. 5C). By contrast, a decrease of *ν*_B_ was observed when Lum or Prelp were targeted to the injury ECM (control^Lum^: (5.324 ± 0.004) GHz, Lum: (5.298 ± 0.006) GHz; control^Prelp^: (5.321 ± 0.005) GHz, Prelp: (5.295 ± 0.005) GHz; Fig. 5C). This suggests that Lum and Prelp increase the tissue compressibility of the local microenvironment in the spinal lesion site.

*ν*_B_ depends not only on *M’* but also on the refractive index (*n*) and density (*ρ*) of the sample in the focal volume (Fig. S9A). To exclude the possibility that the observed changes in *ν*_B_ are solely due to differences in *n*, we quantified its local distribution in the lesion site *in vivo* using optical diffraction tomography (ODT) (Fig. S9B). This revealed 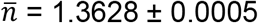 in *pdgfrb*:lum transgenics, and 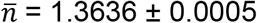 in control animals. In *pdgfrb*:prelp transgenics, we measured 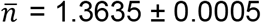 and 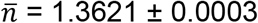 for their respective control animals. To estimate the impact of the uncertainties in 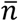 and 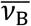 on *M*’ in our experimental framework, we employed Gaussian propagation of uncertainty (Fig. S9C). This showed that the uncertainty in *n* is overall contributing less to the propagated uncertainty in *M*’ than the one of *ν*_B_ for the values reported above. Furthermore, we found that a relative change in *n* contributes less to a relative change in *M*’ than a relative change in *ν*_B_ (Fig. S9D). We conclude that the differences in *M*’ between control and *pdgfrb*:SLRP transgenic animals stem primarily from the observed changes in *ν*_B_ rather than *n*, including respective uncertainties. Hence, the longitudinal moduli with respective propagated uncertainties of *n*, *ν*_B_, refractive index increment of the dry mass content (*α*), and the partial specific volume of the dry mass content (θ) for the different conditionals are calculated to be: *M*’(control^Lum^) = (2.396 ± 0.006) GPa, *M*’(Lum) = (2.373 ± 0.007) GPa, *M*’(control^Prelp^) = (2.394 ± 0.006) GPa, and *M*’(Prelp) = (2.370 ± 0.006) GPa.

Thus, targeting Lum or Prelp to the injury ECM leads to changes in the mechanical properties of the lesion environment and inhibits axon regeneration after SCI.

## DISCUSSION

Zebrafish regenerate severed axons and recover function after CNS injury, in stark contrast to mammals including humans. Here, we demonstrate that differences in injury-associated ECM deposits account for the high regenerative capacity of zebrafish. We show that zebrafish establish a favourable composition of the injury ECM, which is characterized by the low abundance of inhibitory molecules rather than the presence of species-specific axon growth-promoting factors. Our study characterizes the SLRPs Chad, Fmod, Lum, and Prelp as such inhibitory molecules that govern the permissiveness of the injury ECM for regeneration. Increasing the abundance of any one of these proteins in the zebrafish injury ECM is sufficient to impair the capacity of severed axons to regrow across CNS lesions. This identifies SLRPs as previously unknown potent inhibitors of CNS axon regeneration *in vivo*. Furthermore, we demonstrate that CHAD, FMOD, LUM, and PRELP are enriched in the injured human brain and spinal cord. Reducing the abundance of these SLRPs in the injury ECM or attenuating their activity may therefore offer a therapeutic strategy to enhance the permissiveness of CNS lesions for axon growth.

So far, much attention has been focused on constituents of the injury ECM that inhibit regeneration through direct interactions with the axonal growth cone (*4, 5*). However, mechanisms beyond growth cone collapse, repellence, and entrapment are emerging. For example, CSPGs impair inflammation resolution, thereby contributing to a lesion environment hostile to axon growth (*3*). Here, we propose a new class of inhibitors of CNS axon regeneration which function – at least in part – through modifying the structural and mechanical properties of the lesion environment. Our functional experiments support that SLRPs neither inhibit regeneration through i) direct interactions with the axonal growth cone, nor ii) prevention of inflammation resolution, nor iii) altering the composition of the fibroblast-derived injury ECM. Although we cannot exclude that SLRPs influence the availability of neurotrophic factors in the lesion environment, inducing *chad, fmod, lum*, or *prelp* expression specifically in neurons did not affect axonal regrowth *in vivo*, providing evidence against this scenario. Utilizing intravital cross-polarized optical coherence tomography and Brillouin microscopy, we show that the presence of individual SLRP family members in the injury ECM leads to changes in the structural (Fmod, Lum, Prelp) and mechanical (Lum, Prelp; reduced longitudinal modulus) properties of the lesion environment, which coincides with an impaired regenerative capacity of axons. This indicates that a single ECM factor can direct CNS repair toward inhibitory scarring by altering the structural and mechanical properties of the local microenvironment.

Previous studies have shown that CNS injuries in mammals, including humans, are accompanied by a softening of the tissue, whereas the spinal cord of zebrafish stiffens (increased apparent Young’s modulus) (*9, 11, 44*). This work provides *in vivo* evidence for a direct relationship between tissue mechanics (longitudinal modulus) and regenerative outcome upon CNS injury. Furthermore, we propose LUM and PRELP as causal factors of the differential response in zebrafish and humans. LUM has previously been implicated in the modulation of local tissue stiffness during the induction of fold formation in the developing human neocortex, further supporting a role of SLRPs in CNS tissue mechanics (*45*). Interestingly, using Brillouin microscopy, we found that targeting Chad or Fmod to the injury ECM yielded no effective change of the longitudinal modulus, despite their potency in inhibiting axon regeneration being comparable to Lum and Prelp. Whether other mechanical tissue properties, such as the Young’s modulus, are influenced by Chad, Fmod, Lum, or Prelp awaits further investigation.

In conclusion, our data establish the composition, structural organization, and mechanical properties of the ECM as critical determinants of regenerative success after CNS injury. These findings reveal targets to make CNS lesions more conducive to axon growth in mammals, in which regeneration fails.

## ACKNOWLEDGEMENTS

The authors gratefully acknowledge the International Spinal Cord Injury Biobank (ISCIB) for generously providing the human spinal cord specimens used in this project and Dr. Johann Helmut Brandstätter (Biology Department, FAU Erlangen-Nürnberg) for providing mice. We thank Casandra Cecilia Carrillo Mendez and Olga Stelmakh for excellent fish care, Drs Vasiliki Tsata, Jona Kayser, and Catherine Xu for comments on the manuscript. The authors acknowledge financial support from the Else Kröner-Fresenius-Stiftung (K.Ko.), the Deutsche Forschungsgemeinschaft (FR2758/3-1 to R.F.; GU612/8-1 to J.G.), and the Max Planck Society (to J.G.)

## AUTHOR CONTRIBUTIONS

(CRediT nomenclature)

Conceptualization: D.W.; Formal analysis: J.K., K.K., A.P., C.M., G.S., K.S., D.W.; Investigation: J.K., N.J., K.K., C.M., S.M., G.R., V.K., A.P., G.S., T.B., R.F., A.W., N. K., B.S., K.S., K.Ko., D.W.; Methodology: J.K., K.K., S.M., A.W., N.K., B.S., K.S., D.W.; Project administration: D.W.; Resources: I.B., B.S., K.S., J.G., K.Ko. D.W.; Software: K.K., P.M., R.S.; Supervision: K.Ko., D.W.; Visualization: J.K., D.W.; Writing – original draft: J.K., D.W.; Writing – review & editing: J.K., N.J., K.K., C.M., T.B., J.G., K.Ko., D.W.

## DATA AND MATERIALS AVAILABILITY

Except for the proteomics data, all data are available in the main text or the supplementary materials. The shotgun MS data have been deposited to the ProteomeXchange Consortium (http://proteomecentral.proteomexchange.org) via the PRIDE partner repository with the dataset identifier PXD037605 and PXD037590.

## MATERIAL AND METHODS

### Human tissue collection and ethical compliance

This study was approved by the Ethics Committee of the Friedrich-Alexander-University (FAU) Erlangen-Nürnberg, Germany (Refs.#18-193_1-Bio, 193_18B, 92_14B; brain samples) and the Ethics Council of the Max Planck Society (Ref.#2021_40; spinal cord samples) and was conducted in accordance with the Declaration of Helsinki.

For brain samples, informed and written consent was obtained from all patients, their parents, or legal representatives if underage. We reviewed clinical and histological data of individuals who underwent surgery for the treatment of their focal drug-resistant epilepsy and were diagnosed with a scar, i.e., extensively transformed fibrotic and gliotic brain tissue. *En bloc* resections were carried out and tissue was dissected into 5 mm-thick slices along the anterior-posterior axis. Tissue samples were fixed overnight in 4% formalin and routinely processed into liquid paraffin. Six patients with histologically proven scarred brain tissue following traumatic brain injury (TBI, n=3), or repeated surgery (re-OP, n=3) were selected for further investigation. Furthermore, a reference group including no-seizure autopsy controls (n=2) and focal epilepsy patients with a cortical malformation as the primary lesion but no fibrous scarring of resected tissue upon visual inspection (n=4) was also analyzed. In addition to routine hematoxylin and eosin (H&E) staining, immunohistochemical examination of all surgical brain specimens was performed using the following panel of antibodies: mouse monoclonal anti-NeuN (clone A-60, Millipore Cat#MAB377), mouse monoclonal anti-GFAP (clone 6F2, Dako Cat#M0761), and recombinant rabbit monoclonal anti-LUM (Lumican, Invitrogen Cat#MA5-29402). The samples were subsequently digitized using a NanoZoomer Hamamatsu S60 digital slide scanner.

Human spinal cord injury (SCI) samples, control samples, and related clinical and neuropathological information were obtained from the International Spinal Cord Injury Biobank (ISCIB; Vancouver, Canada). The Clinical Research Ethics Board of the University of Columbia (Vancouver, Canada) granted the permission for post-mortem spinal cord acquisition and for sharing biospecimens. Spinal cord biospecimens were collected from consented participants or their next-of-kin and provided as paraffin-embedded tissue sections.

Supplementary Tables S1 and S2 detail clinical and neuropathological data of subjects included in this study.

### Zebrafish husbandry and transgenic lines

All zebrafish lines were kept and raised under a 14/10 h light-dark cycle as described (*46*) according to FELASA recommendations (*47*). We used AB and WIK wild-type strains of zebrafish (*Danio rerio*) and the following transgenic zebrafish lines: BAC(*pdgfrb*:Gal4ff)^ncv24^ (*48*), *UAS*:EGFP^zf82^ (*49*), *UAS-E1b*:Eco.NfsB-mCherry^c264^ (*50*), *pdgfrb*:TetA AmCyan^mps7^ (*12*), *Xla.Tubb*:TetA AmCyan^ue103^ (*13*), and *TetRE*:lum-mCherry^mps3^ (*12*). *elavl3*:GFP-F^mps10^, *TetRE*:aspn-mCherry^mps11^, *TetRE*:chad-mCherry^mps12^, *TetRE*:fmoda-mCherry^mps13^, and *TetRE*:prelp-mCherry^mps14^ transgenic zebrafish lines were established using the DNA constructs and methodology described below.

Combinations of different transgenic zebrafish lines used in this study were abbreviated as follows:

*pdgfrb*:aspn (*pdgfrb*:TetA AmCyan;*TetRE*:aspn-mCherry),
*pdgfrb*:chad (*pdgfrb*:TetA AmCyan;*TetRE*:chad-mCherry),
*pdgfrb*:fmoda (*pdgfrb*:TetA AmCyan;*TetRE*:fmoda-mCherry),
*pdgfrb*:lum (*pdgfrb*:TetA AmCyan;*TetRE*:lum-mCherry),
*pdgfrb*:prelp (*pdgfrb*:TetA AmCyan;*TetRE*:prelp-mCherry),
*Xla.Tubb*:aspn (*Xla.Tubb*:TetA AmCyan;*TetRE*:aspn-mCherry),
*Xla.Tubb*:chad (*Xla.Tubb*:TetA AmCyan;*TetRE*:chad-mCherry),
*Xla.Tubb*:fmoda (*Xla.Tubb*:TetA AmCyan;*TetRE*:fmoda-mCherry),
*Xla.Tubb*:lum (*Xla.Tubb*:TetA AmCyan;*TetRE*:lum-mCherry),
*Xla.Tubb*:prelp (*Xla.Tubb*:TetA AmCyan;*TetRE*:prelp-mCherry),
*pdgfrb*:GFP (BAC(*pdgfrb*:Gal4ff);*UAS*:EGFP),
*pdgfrb*:NTR-mCherry (BAC(*pdgfrb*:Gal4ff);*UAS-E1b*:Eco.NfsB-mCherry).

For live microscopy, embryos were treated with 0.00375% 1-phenyl-2-thiourea (PTU, Sigma-Aldrich Cat#P7629), beginning at 24 hpf (hours post-fertilization), to prevent pigmentation.

All animal experimental procedures were in accordance with institutional and internationally recognized guidelines and were approved by the Regierung von Unterfranken (Government of Lower Franconia, Würzburg, Germany) to comply with German animal protection law. The reference number of the animal experimental permit is RUF 55.2.2-2532.2-1120-15.

### Generation of transgenic zebrafish lines

All primer sequences for molecular cloning are given in Supplementary Data S1. To create the donor plasmid for *elavl3*:GFP-F transgenic zebrafish, the sequence coding for the membrane-localized GFP (EGFP fused to farnesylation signal from c-HA-Ras) was amplified from the pEGFP-F vector (Clonetech) using primer pair #1, and cloned downstream of the zebrafish *elavl3* promoter (Addgene plasmid Cat#59530) (*51*). To create the donor plasmids for generation of *TetRE*:aspn-mCherry, *TetRE*:chad-mCherry, *TetRE*:fmoda-mCherry, and *TetRE*:prelp-mCherry transgenic fish, the sequences coding for zebrafish *aspn* (ENSDART00000064798.6), *chad* (ENSDART00000066264.4), *fmoda* (ENSDART00000065985.5), and *prelp* (ENSDART00000155521.2) were amplified from cDNA of developing zebrafish (primer pairs #2-5), fused to mCherry at the C-terminus and cloned downstream of the tetracycline operator sequence as described (*52, 53*). Generation of *elavl3*:GFP-F, *TetRE*:aspn-mCherry, *TetRE*:chad-mCherry, *TetRE*:fmoda-mCherry, and *TetRE*:prelp-mCherry transgenic zebrafish lines was achieved by injection of 35 pg of the respective donor plasmid together with *in vitro* synthesized capped sense mRNA of the Tol2 transposase into 1-cell embryos (*54*).

### Drug treatments

Drug treatments were performed according to the schematic timelines shown with each experiment. Doxycycline (DOX; Sigma-Aldrich Cat#D9891) was dissolved in reverse osmosis H_2_O at 50 mg/mL and used at a final concentration of 25 μg/mL. Metronidazole (MTZ; Sigma-Aldrich #M3761) was dissolved in DMSO at 800 mM stock concentration and used at a final concentration of 2 mM.

### Zebrafish spinal cord lesions and behavioral recovery

A detailed protocol for inducing spinal cord lesions in zebrafish larvae has been previously described (*55*). Briefly, zebrafish larvae (3 dpf) were anesthetized in E3 medium containing 0.02% MS-222. A 30 G x ½” hypodermic needle was used to transect the spinal cord by either incision or perforation at the level of the urogenital pore. After surgery, larvae were returned to E3 medium for recovery and kept at 28.5°C. Larvae that had undergone extensive damage to the notochord were excluded from further analysis. For lesions, the experimenter was blinded to the experimental treatment. Larvae used for lesions were randomly taken from Petri dishes containing up to 50 animals, however, no formal randomization method was used.

Analysis of behavioral recovery after spinal cord transection in larval zebrafish was performed as previously described (*17*), using EthoVision XT software (Noldus). Behavioral data are shown as the distance traveled within 10 s after touch, averaged for triplicate measures per larvae.

### Image acquisition and processing

Images were acquired using the systems described in each subsection. Images were processed using ImageJ (http://rsb.info.nih.gov/ij/), Adobe Photoshop CC, and Zeiss ZEN blue software. Figures were assembled using Adobe Photoshop CC.

### Live imaging of larval zebrafish

For live confocal imaging, zebrafish larvae were anesthetized in E3 medium containing 0.02% MS-222 and mounted in the appropriate orientation in 1% low melting point agarose (Ultra-Pure^™^ Low Melting Point, Invitrogen Cat#16520) between two microscope cover glasses. During imaging, larvae were covered with 0.01% MS-222-containing E3 medium to keep preparations from drying out. Imaging was done using a Plan-Apochromat 10x/0.45 M27 objective, Plan-Apochromat 20x/0.8 objective, and C-Apochromat 40x/1.2 W Korr UV-VIS-IR objective on a Zeiss LSM 980 confocal microscope.

### Sectioning of larval zebrafish

Terminally anesthetized zebrafish larvae were fixed in 4% paraformaldehyde (PFA; Thermo Fisher Scientific Cat#28908) in PBS for 1 h at room temperature. After two washes in PBT (0.1% Tween-20 in PBS), larvae were embedded in 4% agarose in PBS and 50-100 μm sections were obtained using a vibratome (Leica, VT1200S). Sections were counterstained with DAPI (Thermo Fisher Scientific Cat#62248) to visualize nuclei and mounted in 75% glycerol. Images were acquired using a Plan-Apochromat 20x/0.8 on a Zeiss LSM 980 confocal microscope.

### Tissue dissociation and FACS

GFP^+^ and GFP^-^ cells were isolated from *pdgfrb:GFP* transgenic zebrafish at 4 dpf. 120 animals were pooled for each experiment. Before tissue dissociation, head, yolk sac, and tail fin were removed, using micro scissors. Dissociation, live cell staining, and FACS sorting of cells was done as previously described (*12*). Briefly, excised tissue was enzymatically dissociated using 0.25% Trypsin-EDTA (Gibco Cat#25200072) followed by mechanical dissociation using a fire-polished glass Pasteur pipette. Cell suspension was filtered through a 20 μm cell strainer (pluriSelect Cat#43-10020-40). Following centrifugation for 10 min at 300 g, cells were resuspended in 500 μL HBSS medium (Gibco Cat#14065-056). To stain for viable cells, Calcein Blue (Invitrogen Cat#C1429) was added to the cell suspension. Cells were directly sorted into lysis buffer using a MoFlo Astrios EQ sorter (Beckman Coulter). For the detection of GFP, a 488 nm excitation laser and a 526/52 nm bandpass filter were used. Calcein Blue was detected following 405 nm excitation using a 431/28 nm bandpass emission filter.

### Quantitative RT-PCR (qRT-PCR)

For qRT-PCR on isolated *pdgfrb*^+^ cells, total RNA was extracted from ~70,000 FACS-sorted GFP^+^ cells of dissected trunks from unlesioned *pdgfrb*:GFP transgenic animals at 4 dpf, using Total RNA Purification Plus Micro Kit (Norgen Biotek Cat#48500). The total RNA extracted from an equal number of GFP^-^ cells obtained from the same animals served as control.

For quantification of transcript levels in the lesion site at 1 dpl, trunk tissue of 75 animals spanning approximately three somites in length and containing the lesion site were isolated using micro scissors. Corresponding trunk tissue of unlesioned age-matched clutch mates served as control. Total RNA was extracted using TRIzol reagent (Invitrogen Cat#15596026).

Reverse transcription was performed with Maxima^™^ H Minus cDNA Synthesis Master Mix with dsDNase (Thermo Fisher Scientific Cat#1681), using a combination of oligo(dT) and random hexamer primers. qRT-PCR was performed at 60°C using PowerUp^™^ SYBR^™^ Green Master Mix (Applied Biosystems Cat#A25918) on a StepOnePlus^™^ Real-Time PCR system. Samples were run in triplicates, and expression levels were normalized to *actb1* control. Normalized relative mRNA levels were determined using the ΔΔCt method.

Primers were designed to span an exon–exon junction using Primer-BLAST software (https://ncbi.nlm.nih.gov/tools/primer-blast/). All primer sequences are given in Supplementary Data S1.

### Sample preparation for mass spectrometry (MS) analysis

For label-free MS-based quantitative proteomics, proteins were isolated from trunk tissue of 75 larvae spanning approximately three somites in length and containing the lesion site or corresponding unlesioned age-matched trunk tissue. To identify injury-induced changes in protein abundance, MS analysis was performed on three biological replicates for each condition. To identify changes in protein abundance caused by targeting SLRPs to *pdgfrb*^+^ cells, MS analysis was performed on four biological replicates for each condition. Isolation of proteins was done as described in (*56*) with minor modifications. Proteins were extracted in three distinct fractions, and each fraction was analyzed separately by LC-MS/MS. Zebrafish tissue was homogenized in PBS for 10x 30 s at high intensity, and 10x 30 s pauses in between, using the Bioruptor Plus sonication system (Diogenode Cat#UCD-300). After centrifugation at 16,000 g for 2 min, the supernatant containing the soluble proteins was collected (fraction 1). The pellet was resuspended in detergent-containing buffer (50 mM Tris (Sigma-Aldrich Cat#T1503), 5% glycerin (Roth Cat#4043.2), 500 mM NaCl (VWR Cat#27810.295), 1% NP40 (Fluka Cat#74385), 2% SDC (Sigma-Aldrich Cat#SER0046), 1% SDS (Roth Cat#4360.2), 1% DNAse I (in House made, ProteinProduction Facility), 1 mM MgCl_2_ (Sigma-Aldrich Cat#M9272)), followed by incubation at 0°C for 20 min and homogenization using the Bioruptor Plus sonication system. After centrifugation at 16,000 g for 2 min, the supernatant containing the detergent-soluble proteins (fraction 2) and the detergent-insoluble protein pellet (fraction 3) were collected. Proteins of fraction 1 and fraction 2 were precipitated by incubation with acetone at −20°C for overnight. The pellets of all three fractions were dissolved in reduction and alkylation buffer (6M guanidinium chloride (Merck Cat#1.04220.1000), 100 mM Tris-HCl pH 8.5, 10 mM TCEP (Thermo Fisher Scientific Cat#77720), 50 mM CAA (Sigma-Aldrich Cat#C0267)), followed by incubation at 99°C for 15 min and sonication in the Bioruptor Plus sonication system. The proteins were diluted 1:3 with Urea-buffer (4.5 mM Urea (Sigma-Aldrich Cat#U1250), 10 mM Tris-HCl, 3% Acetonitrile (VWR Cat#83640.320), 1 μg of LysC (Wako Cat#129-02541), and incubated at 37°C for 3 h. Thereafter, the digestion mixture was diluted 1:3 with 10% Acetonitrile in MS grade water (VWR Cat#83645.320), followed by sonication in the Bioruptor Plus sonication system and incubation with 1 μg of LysC and 2 μg of Trypsin (Promega Cat#V511A) at 37°C for overnight. Acetonitrile was removed using a SpeedVac (Christ Cat#RVC 2-25 together with CT 02-50), and peptides were further purified using in-house produced three plug SCX stage tips (Empore^™^ Cation Solid Phase Extraktion Disks Cat#2251). After elution with 60 μL of 5% ammonia solution (Roth Cat#5460.1) in 80% acetonitrile, samples were vacuum-dried in the SpeedVac.

### LC-MS/MS acquisition

Peptides were solubilized in 6 μL buffer A (100% MS-LC grade water and 0.1% Formic acid (VWR Cat#84865.180), and a total volume of 3 μL were loaded onto a 30 cm-long column (75 μm inner diameter (Polymicro Cat#TSP075375); packed in-house with ReproSil-Pur 120 C18-AQ 1.9-micron beads (Dr. Maisch GmbH Cat#r119.aq)) via the autosampler of the Thermo Scientific Easy-nLC 1200 (Thermo Fisher Scientific) at 60°C. For the identification of injury-induced changes in protein abundance (samples: unlesioned 4 dpf, 1 dpl; unlesioned 5 dpf, 2 dpl), eluted peptides were directly sprayed onto the Q Exactive HF Orbitrap LC-MS/MS system (Thermo Fisher Scientific), using the nanoelectrospray interface. As a gradient, the following steps were programmed with increasing addition of buffer B (80% acetonitrile, 0.1% formic acid): linear increase from 30% over 120 min, followed by a linear increase to 60% over 10 min, followed by a linear increase to 95% over the next 5 min, and finally buffer B was maintained at 95% for another 5 min. The mass spectrometer was operated in a data-dependent mode with survey scans from 300 to 1750 m/z (resolution of 60000 at m/z = 200), and up to 15 of the top precursors were selected and fragmented using higher energy collisional dissociation (HCD with a normalized collision energy of value of 28). The MS2 spectra were recorded at a resolution of 15k (at m/z = 200). AGC target for MS1 and MS2 scans were set to 3E6 and 1E5, respectively, within a maximum injection time of 100 ms for MS1 and 25 ms for MS2. Dynamic exclusion was set to 16 ms. For identification of changes in protein abundance caused by overexpression of SLRPs, eluting peptides were directly sprayed onto the timsTOF Pro LC-MS/MS system (Bruker). As gradient, the following steps were programmed with increasing addition of buffer B (80% acetonitrile, 0.1% formic acid): linear increase from 5% to 25% over 75 min, followed by a linear increase to 35% over 30 min, followed by a linear increase to 58% over the next 5 min, followed by a linear increase to 95% over the next 5 min, and finally buffer B was maintained at 95% for another 5 min. Data acquisition on the timsTOF Pro was performed using timsControl. The mass spectrometer was operated in data-dependent PASEF mode with one survey TIMS-MS and ten PASEF MS/MS scans per acquisition cycle. Analysis was performed in a mass scan range from 100-1700 m/z and an ion mobility range from 1/K0 = 0.85 Vs cm^-2^ to 1.30 Vs cm^-2^, using equal ion accumulation and ramp time in the dual TIMS analyzer of 100 ms each at a spectra rate of 9.43 Hz. Suitable precursor ions for MS/MS analysis were isolated in a window of 2 Th for m/z < 700 and 3 Th for m/z > 700 by rapidly switching the quadrupole position in sync with the elution of precursors from the TIMS device. The collision energy was lowered as a function of ion mobility, starting from 59 eV for 1/K0 = 1.6 Vs cm^-2^ to 20 eV for 0.6 Vs cm^-2^. Collision energies were interpolated linearly between these two 1/K0 values and kept constant above or below these base points. Singly charged precursor ions were excluded with a polygon filter mask and further m/z and ion mobility information was used for ‘dynamic exclusion’ to avoid re-sequencing of precursors that reached a ‘target value’ of 20,000 a.u. The ion mobility dimension was calibrated linearly using three ions from the Agilent ESI LC/MS tuning mix (m/z, 1/K0: 622.0289, 0.9848 Vs cm^-2^; 922.0097 Vs cm^-2^, 1.1895 Vs cm^-2^; 1221.9906 Vs cm^-2^, 1.3820 Vs cm^-2^).

### Computational MS data analysis

Raw data were processed using the MaxQuant computational platform (versions 1.6.17.0 and 2.0.1.0) (*57*) with standard settings applied. Briefly, the peak list was searched against the zebrafish (*Danio rerio*) proteome database (SwissProt and TrEMBL, 46847 entries) with an allowed precursor mass deviation of 4.5 ppm and an allowed fragment mass deviation of 20 ppm. MaxQuant enables individual peptide mass tolerances by default, which was used in the search. Cysteine carbamidomethylation was set as static modification, and methionine oxidation and N-terminal acetylation as variable modifications. Proteins were quantified across samples using the label-free quantification algorithm in MaxQuant, generating label-free quantification (LFQ) intensities. The match-between-runs option was enabled.

### Bioinformatic analysis of proteomics data

LFQ intensity values were imported to Perseus software (version 1.6.14.0) (*58*). Filters were set to exclude proteins identified by site, matching to the reverse database or contaminants. For identification of injury-induced changes in protein abundance, proteins that did not exhibit valid values for all three replicates of at least one experimental condition (unlesioned 4 dpf, 1 dpl; unlesioned 5 dpf, 2 dpl) were excluded. This was necessary to include proteins undetectable in unlesioned animals but with enrichment after SCI. For identification of changes in protein abundance caused by cell-type specific induction of SLRPs, all proteins that exhibited ≥ 1 invalid value in all eight samples were excluded. Proteins exhibiting significantly altered abundance between experimental and control samples were identified by permutation-based analysis (*59*) with FDR < 0.1, s0 = 0.1, and |FC| ≥ 1.3. Principal component analysis plots and volcano plots were created using Perseus software.

### Reactome pathway analysis

To identify overrepresented biological pathways based on the observed protein abundance ratios, Reactome pathway analysis was performed using g:Profiler (*60, 61*). Proteins exhibiting significantly increased abundance in lesioned samples as compared to corresponding unlesioned age-matched control samples (FDR < 0.1, FC ≥ 1.3, s_0_ = 0.1) were analyzed. Reactome pathways with an adjusted (Bonferroni) *P*-value ≤ 0.05 were considered significantly enriched.

### Heatmaps

The zebrafish matrisome was obtained from http://matrisome.org (*18*) and manually updated for missing UniProt identifiers. The updated matrisome list was imported to Perseus software and matched with the MS dataset to add missing UniProt identifiers, gene symbols, as well as further specifications, including division (core matrisome, matrisome-associated, and putative matrisome proteins) and protein symbol of mammalian orthologues. The list of matrisome proteins exhibiting significantly altered abundance between lesioned and corresponding unlesioned age-matched control samples (FDR < 0.1, |FC| ≥ 1.3, s_0_ = 0.1) was extracted, and the Z-score was calculated. Heatmaps were created using Perseus software.

### Cross-species comparison of proteomics data

For cross-species comparison of proteomics data, a published dataset from adult Sprague-Dawley rats at seven days post-contusion spinal cord injury (T10 injury level) and uninjured control T10 spinal cord segments was re-analyzed (*30, 62*). LFQ intensity values of ECM-enriched 4M guanidine spinal cord tissue extracts (https://data.mendeley.com/datasets/npkwh5vsss/1) were imported to Perseus software. Filters were set to exclude proteins identified by site, matching to the reverse database or contaminants. Additionally, proteins were excluded that did not exhibit valid values for all three replicates of at least one experimental condition (unlesioned, 7 dpl). Proteins exhibiting significantly altered abundance between lesioned and unlesioned control samples were identified by permutation-based analysis (*59*) with FDR < 0.1, s_0_ = 0.1, and |FC| ≥ 1.3. A pairwise comparison of the obtained protein lists using Venny 2.1 software (https://bioinfogp.cnb.csic.es/tools/venny) was performed to identify differentially regulated matrisome proteins between rat and zebrafish (*63*).

### In situ hybridization (ISH)

A detailed protocol for ISH on whole-mount zebrafish larvae with digoxygenin (DIG)-labeled antisense probes has been described in (*55*). In brief, terminally anesthetized larvae were fixed in 4% PFA (Thermo Fisher Scientific Cat#28908) in PBS and treated with Proteinase K (Invitrogen Cat#25530-049) followed by re-fixation for 15 min in 4% PFA in PBS. DIG-labeled antisense probes were hybridized overnight at 65°C. This protocol allows efficient probe penetration in whole-mount preparations of 5 dpf larvae (*13*). Color reaction was performed after incubation with anti-DIG antibody conjugated to alkaline phosphatase (Sigma-Aldrich Cat#11093274910) using NBT/BCIP substrate (Roche Cat#11697471001). Samples were mounted in 75% glycerol in PBS and imaged in multi-focus mode using a Leica M205 FCA stereo microscope equipped with a Leica DMC6200 C color camera.

Information on ISH probes, including primer sequences used for molecular cloning, is provided in Supplementary Data S1. All ISH probes showed specific staining in developmental domains.

### Immunofluorescence (IF)

A detailed protocol for IF on whole-mount zebrafish larvae has been previously described (*55*). Briefly, terminally anesthetized larvae were fixed in 4% PFA (Thermo Fisher Scientific Cat#28908) in PBS for 1 h at room temperature. After removing the head and tail using micro scissors, larvae were permeabilized by subsequent incubation in acetone and Proteinase K (Invitrogen Cat#25530-049). Samples were re-fixed in 4% PFA in PBS, blocked in PBS containing 1% Triton X-100 (PBTx) and 4% bovine serum albumin, and incubated over two to three nights with the primary antibody of interest. After several washes in PBTx, samples were incubated over two nights with the secondary antibody of interest. Thereafter, samples were washed in PBTx and mounted in 75% glycerol in PBS. Imaging was performed using a Plan-Apochromat 20x/0.8 objective on a Zeiss LSM 980 confocal microscope.

For neurite outgrowth assay, dorsal root ganglion (DRG) neuron cultures were fixed in 4% PFA (Thermo Fisher Scientific Cat#28908) in PBS for 15 min at room temperature. After permeabilization with 0.1% Triton X-100 in PBS for 20 min at 37°C, DRG neurons were blocked with 5% goat serum (Sigma-Aldrich Cat#G9023) in PBS for 45 min at 37°C. Neurites were labeled by subsequent incubation with anti-Tubulin β3 antibody in PBS for 2 h and fluorophore-conjugated secondary antibody in PBS for 1.5 h at 37°C. DRG neurons were mounted in Fluoromount-G with DAPI (Invitrogen Cat#E132139) and imaged using a Plan-Apochromat 20x/0.8 objective and digital camera system on a Zeiss LSM 980 microscope.

IF on human paraffin-embedded tissue sections was performed as previously described (*64*). Imaging was performed using Plan-Apochromat 10x/0.45 M27 and Plan-Apochromat 20x/0.8 objectives on a Zeiss LSM 980 confocal microscope.

We used rabbit polyclonal anti-Collagen IV (Abcam Cat#ab6586), mouse monoclonal anti-Tubulin β3 (TUBB3, Biolegend Cat#801201), rabbit polyclonal anti-Mpx (GeneTex Cat#GTX128379), mouse monoclonal anti-GFAP (clone GA5, Cell Signaling Technology Cat#mAB3670), recombinant rabbit polyclonal anti-GFAP (clone RM1003, Abcam Cat#ab278054), rabbit monoclonal anti-LUM (Invitrogen Cat#MA5-29402), sheep polyclonal anti-PRELP (R&D Systems Cat#AF6447), rabbit polyclonal anti-CHAD (Invitrogen Cat#PA-553761), mouse monoclonal anti-CHAD (clone 8B7, Thermo Fisher Scientific Cat#H00001101-M01), and mouse monoclonal anti-FMOD (clone 549302, R&D Systems Cat#MAB5945) primary antibodies. Secondary fluorophore-conjugated antibodies were from Invitrogen.

### Whole-mount TUNEL/anti-GFP co-labeling

Terminally anesthetized larvae were fixed in 4% PFA (Thermo Fisher Scientific Cat#28908) in PBS for overnight at 4°C. After removing the head and tail using micro scissors, larvae were permeabilized by subsequent incubation in acetone and Proteinase K (Invitrogen Cat#25530-049) as described for whole-mount IF. Samples were re-fixed in 4% PFA in PBS and Click-iT TUNEL Alexa Fluor 647 Imaging Assay (Thermo Fisher Cat#C10247) was performed according to the manufacturer’s protocol to label apoptotic cells. Briefly, samples were equilibrated in TdT reaction buffer for 30 min at room temperature, followed by incubation in TdT reaction cocktail for overnight at room temperature. The click-it reaction was performed for 60 min at room temperature. Thereafter, samples were blocked in PBTx (1% Triton X-100 in PBS) containing 3% bovine serum albumin and incubated over three nights with chicken anti-GFP antibody (Abcam Cat#ab13790). After several washes in PBTx, samples were incubated over two to three nights with the secondary antibody of interest. Thereafter, samples were washed in PBTx and mounted in 75% glycerol in PBS. Imaging was done using a Plan-Apochromat 10x/0.45 M27 objective on a Zeiss LSM 980 confocal microscope.

### Primary dorsal root ganglia (DRG) culture/neurite outgrowth assay

Handling of mice was performed in accordance with animal welfare laws, complied with ethical guidelines, and was approved by the responsible local committees and government bodies (University of Erlangen, Amt für Veterinärwesen der Stadt Erlangen, and the Regierung von Unterfranken). Adult (4-6 months old) C57BL/6J mice were killed by cervical dislocation, and spinal cords were removed. DRG neurons were dissected from the spinal cord, incubated in Neurobasal medium (NB, Thermo Fisher Scientific Cat#21103049) containing 2.5 mg/mL collagenase P (Sigma-Aldrich Cat#11213857001) and maintained in an incubator for 1 h at 37°C in 5% CO_2_. The DRG tissue was homogenized by pipetting using finely fired-polished drawn glass pipettes. Dissociated DRG neurons were separated from axon stumps and myelin debris via a 14% bovine serum albumin layer (Sigma-Aldrich Cat#A9205) and centrifugation at 120 rpm for 8 min. The pellet with DRG neurons was resuspended in NB containing 20 μL/mL B27 supplement 50x (Gibco Cat#17504044), 2 mM Glutamax (ThermoFisher Cat#35050-061), 10 μM/mL antibiotic-antimycotic (Gibco Cat#15240062), and 0.01 μg/mL nerve growth factor (ThermoFisher Cat#3257-019). Isolated DRG neurons were seeded on glass coverslips coated with 0.1 mg/mL Poly-D-lysine (PDL, Gibco Cat#A3890401), followed by coating with either 10 μg/mL Laminin (ThermoFisher Cat#23017015), a mix of 10 μg/mL Laminin and 5 μg/mL of each SLRP protein (recombinant human CHAD (R&D Systems Cat#8218-CH), FMOD (R&D Systems Cat#9840-FM), LUM (Abcam Cat#ab221400), and PRELP (R&D Systems Cat#6447-PR)), or a mix of 10 μg/mL Laminin and 5 μg/mL CSPGs (Sigma-Aldrich Cat#CC117). All recombinant SLRP proteins used were produced in mammalian expression systems. DRG cultures were incubated for 48 h at 37°C and 5% CO_2_ to allow neurite outgrowth.

### Cross-polarized optical coherence tomography (CP-OCT)

For *in vivo* CP-OCT (*37*), 1 dpl zebrafish larvae were anesthetized in E3 medium containing 0.015% MS-222 and mounted in a lateral position in 1% low gelling temperature agarose (Sigma-Aldrich Cat#A0701) between two microscope cover glasses. During imaging, larvae were covered with 0.01% MS-222-containing E3 medium to keep preparations from drying out. Cross-polarized images of the spinal lesion site were acquired using a custom-built CP-OCT system (*65*). Light from a broadband supercontinuum laser (YSL Photonics Cat#SC-OEM) was filtered to acquire a spectrum centered at 885 nm with a full width at half maximum of 80 nm. The laser was operated at 200 MHz. The filtered spectrum from the laser was coupled to a single-mode optical fiber and collimated using a collimator (Thorlabs Cat#F230APC-850). The light was further split using a 90/10 beam splitter (Thorlabs Cat#BS025) into a reference beam and a sample beam. The sample was illuminated with 15 mW of optical power. A combination of a quarter-wave plate (Thorlabs Cat#SAQWP05M-1700) and a lens (Thorlabs Cat#AC254-030-AB) was inserted in the reference and the sample arm to control the polarization of the light. A galvano mirror (Thorlabs Cat#GVS012) was used to scan the laser beam over the sample. The reflected reference and the sample signals were acquired using a custom-designed spectrometer consisting of a reflective collimator (Thorlabs Cat#RC08APC), a holographic grating (Wasatch Photonics Cat#1200 l/mm@840nm), a lens (Thorlabs Cat#AC-254-080-B), and a line scan camera (Basler Cat#2048 pixels, ral2048-48gm) operating at 25 kHz line scan rate. The spectrometer signal was processed using LabVIEW-based custom-designed software (National Instruments). The acquired interference spectrum from the camera was spectrally recalibrated from wavelength space to wavenumber space (*66*) and a fast Fourier transform was performed to obtain an axial profile of the sample.

### Combined confocal fluorescence and Brillouin microscopy (BM)

For *in vivo* BM, 1 dpl zebrafish larvae were anesthetized in E3 medium containing 0.015% MS-222 and mounted in a lateral position in 1% low gelling temperature agarose on a 35 mm glass-bottom dish (ibidi Cat#81158). During imaging, larvae were covered with 0.015% MS-222-containing E3 medium to keep preparations from drying out. Brillouin frequency shift images were acquired by BM employing a confocal configuration and a Brillouin spectrometer consisting of a two-stage virtually imaged phase array (VIPA) etalon, as previously described in detail elsewhere (*43*). Briefly, the sample was illuminated by a frequency-modulated diode laser beam (*λ* = 780.24 nm, DLC TA PRO 780, Toptica), which was stabilized to the D_2_ transition of rubidium ^85^Rb. A Fabry-Perot interferometer in two-pass configuration and a monochromatic grating (Toptica) were employed to suppress further the contribution of amplified spontaneous emission to the laser spectrum. The laser light was coupled into a single-mode fiber and guided into the backside port of a commercial inverted microscope stand (Axio Observer 7, Zeiss). An objective lens (20x, NA = 0.5, EC Plan-Neofluar, Zeiss) illuminated the sample on a motorized stage with an optical focus. The laser power at the sample plane was set at 15 mW. The backscattered light from the sample was collected by the same objective lens, coupled into the second single-mode fiber to achieve confocality, and delivered to a Brillouin spectrometer. In the Brillouin spectrometer, the backscattered light was collimated and passed through a molecular absorption cell filled with rubidium ^85^Rb (Precision Glassblowing Cat#TG-ABRB-I85-Q), in which the intensity of the Rayleigh scattered and reflected light were significantly suppressed. After passing through the molecular absorption cell, the beam was guided to two VIPA etalons (Light Machinery Cat#OP-6721-6743-4) with the free spectral range of 15.2 GHz, which convert the frequency shift of the light into the angular dispersion in the Brillouin spectrum. The Brillouin spectrum was acquired by a sCMOS camera (Teledyne Cat#Prime BSI), with the exposure time of 0.5 s per measurement point. The two-dimensional Brillouin frequency map of the injured region was measured by scanning the motorized stage on the microscope stand, with the translational step size of 0.5 μm. The Brillouin microscope was controlled with custom acquisition software written in C++ (https://github.com/BrillouinMicroscopy/BrillouinAcquisition). Confocal fluorescence imaging was performed in the same region of interest (ROI) as Brillouin measurement using a re-scan confocal microscopy (RCM) module (RCM2, Confocal.nl) that was attached to one side port of the microscope stand. The RCM module consists of a sCMOS camera (Prime BSI Express, Teledyne) and the multi-line laser unit (Skyra, Cobolt) as an excitation illumination source for four laser lines (*λ* = 405 nm, 488 nm, 562 nm, and 637 nm). A pinhole of the diameter of 50 μm and the re-scanning imaging principle (*67*) provides rapid confocal fluorescence imaging with a lateral resolution of 120 nm. By focusing a confocal fluorescence image in the center of the spinal cord of *elavl3*:GFP-F transgenic zebrafish larvae, the axial plane of Brillouin imaging was determined.

### Optical diffraction tomography (ODT)

For *in vivo* ODT, 1 dpl zebrafish larvae were anesthetized in E3 medium containing 0.015% MS-222 and mounted in a lateral position in 1% low gelling temperature agarose on a 35 mm glass-bottom dish (ibidi Cat#81158). During imaging, larvae were covered with E3 medium containing 0.015% MS-222 and 20% of refractive index (RI)-matching agent Iodaxanol (OptiPrep^™^; Sigma-Aldrich Cat#D1556) (*68*). Iodaxanol was used to reduce the RI difference between the zebrafish larvae and the surrounding medium. The final RI of the medium was 1.351, which was determined by an ABBE refractometer (Kern & Sohn GmbH Cat#ORT1RS). The RI distribution of the zebrafish spinal lesion site was measured by ODT employing Mach-Zehnder interferometry to measure multiple complex optical fields from various incident angles, as previously described (*69*). A solid-state laser beam (λ = 532 nm, 50 mW, CNI Optoelectronics Technology Co.) was split into two paths using a beamsplitter. One beam was used as a reference beam and the other beam illuminated the sample on the stage of an inverted microscope (Axio Observer 7, Carl Zeiss AG) through a tube lens (*f* = 175 mm) and a water-dipping objective lens (40x, NA = 1.0, Carl Zeiss AG). A high numerical aperture objective lens (40x, water immersion, NA = 1.2, Carl Zeiss AG) collected the beam diffracted by the sample. To reconstruct a 3D RI tomogram of the sample, the sample was illuminated from 150 different incident angles scanned by a dual-axis galvano mirror (Thorlabs Cat#GVS212/M) located at the conjugate plane of the sample. The diffracted beam interfered with the reference beam at an image plane and generated a spatially modulated hologram, which was recorded with a CMOS camera (XIMEA Cat#MQ042MG-CM-TG). The field-of-view of the camera covers 205.0 μm × 205.0 μm. The complex optical fields of light scattered by the samples were retrieved from the recorded holograms by applying a Fourier transform-based field retrieval algorithm (*70*). The 3D RI distribution of the samples was reconstructed from the retrieved complex optical fields via the Fourier diffraction theorem, employing the first-order Rytov approximation (*71, 72*). A more detailed description of tomogram reconstruction can be found elsewhere (*73*). The MATLAB script for ODT reconstruction can be found at https://github.com/OpticalDiffractionTomography/ODT_Reconstruction.

### Quantifications and statistics

Unless otherwise indicated, controls refer to DOX-treated clutch mates of wild type, single transgenic Tet-activator or single transgenic Tet-responder genotype.

For quantification of axonal bridge thickness, transgenic live animals (*elavl3*:GFP-F) were imaged using a Plan-Apochromat 20x/0.8 objective on a Zeiss LSM 980 confocal microscope. The length of a vertical line that covers the width of the axonal bridge at the center of the lesion site, was then determined using ImageJ software (https://imagej.nih.gov/ij/index.html). Measurements of axonal bridge thickness were performed in samples of three independent experiments. Spinal cord lesions for one out of the three experimental replicates were induced by a second operator.

Quantification of fluorescence signal in the lesion site was performed on captured images of whole-mount samples using ImageJ software, following previously published protocols (*13, 20*). Confocal image stacks were collapsed before analysis. For the quantification of area coverage of *pdgfrb*:GFP^+^ in the lesion site, collapsed confocal stacks were converted to a binary image using the automated mean of grey levels thresholding function of ImageJ software. A pre-set ROI of constant size was applied to all images of the same experiment and the number of pixels determined (pixel area). The ROI was placed in the center of the lesion site and ventrally limited by the notochord.

Quantification of *pdgfrb*:GFP^+^/TUNEL^+^ cells was performed in a pre-set ROI of constant size on optical sections of a confocal stack using ImageJ software. The ROI was ventrally limited by the dorsal/caudal artery. *pdgfrb*:GFP^+^/TUNEL^+^ cells in the floor plate or median fin fold were excluded from analysis based on their location and morphology. Quantification of *Mpx*^+^ cells in the lesion site was performed in a pre-set ROI of constant size on optical sections of a confocal stack using ImageJ software. The ROI was ventrally limited by the notochord.

The neurite length of adult murine DRG neurons was quantified using the semi-automated SNT Fiji-ImageJ plugin (*74*). The longest neurite of each DRG neuron was identified and measured.

Extraction and quantification of Brillouin frequency shifts were performed in a pre-set ROI, using custom software written in Python (https://github.com/GuckLab/impose; https://github.com/BrillouinMicroscopy/BMicro). The ROI was 40 μm x 20 μm, which corresponds to the determined average axonal bridge thickness and the average distance between the spinal cord stumps following a dorsal incision lesion at 1 dpl. The ROI was placed in the center of the lesion site and ventrally limited 5 μm dorsal to the notochord. The ROI was applied to all images. Brillouin frequency shifts were measured in samples of at least three independent experiments. The measured Brillouin frequency shift *ν_B_* can be expressed in terms of the longitudinal modulus *M*’, refractive index (RI) *n*, and density *ρ* of the specimen, as well as the incident wavelength *λ*_0_ and scattering angle *ϑ* given by the setup: 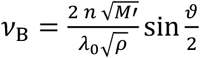. All measurements were performed in the backscattering configuration with *ϑ* = 180° and, accordingly 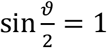. The longitudinal compressibility *κ_L_* may be expressed as the inverse of the longitudinal modulus 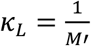. For the sake of conciseness *κ_L_* will be referred to as ‘compressibility’ in the rest of the text, despite the fact that this term is usually defined as the inverse of the bulk modulus *K*’. Bulk modulus *K*’, shear modulus *G*’, and longitudinal modulus *M*’ are related by the following equation in the case of an isotropic sample: 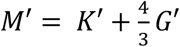.

To quantify the co-polarization ratio (ratio of preserved polarization to total reflectivity) in the lesion site, two images of the tissue were acquired; one co-polarized and one cross-polarized image. For the acquisition of the co-polarized image, the angle of the quarter-wave plate (QWP) axis in the reference path was set in such a way that the light in the sample and the reference arm had the same polarization. To acquire the cross-polarized image, the QWP in the reference beam path was rotated by 45°, resulting in orthogonal polarization between reference and sample arm. Sample reflectivity *R*(*z*) and co-polarization ratio *δ*(*z*) of the sample was calculated using the amplitude of the co-(*A*_co_) and cross-(*A_cross_*) polarized images using the following equations: 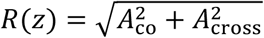 and 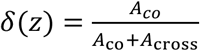. Using ImageJ software, the co-polarization ratio was determined in a pre-set ROI of constant size for the averaged intensity projection of the complete image stack. The ROI was placed in the center of the lesion site and ventrally limited by the notochord. A second independent observer validated the results.

Extraction and quantification of the RI and mass density distribution from reconstructed tomograms were performed using custom-written MATLAB (MathWorks) scripts. The mass density of the samples was calculated directly from the reconstructed RI tomograms, since the RI of the samples, *n*(*x, y, z*), is linearly proportional to the mass density of the material, *α*(*x,y,z*), as *n*(*x,y,z*) = *n_m_* + *αρ*(*x,y,z*), where *n_m_* is the RI value of the surrounding medium and *α* is the RI increment (*dn/dc*), with *α* = 0.1919 mL/g for proteins and nucleic acids (*75, 76*). RI and mass density distribution from reconstructed tomograms were evaluated in a pre-set ROI in a sagittal slice of the RI tomogram along the *x-y* plane at the focused plane, using a custom-written MATLAB (MathWorks) script. The ROI was the same as applied to Brillouin and ODT images. Note that the RI was measured in samples independent of those analyzed with BM.

Unless indicated, no data were excluded from analyses. Except for determining effect sizes and Gaussian propagation of uncertainty, all statistical analyses were performed using Graph Pad Prism 9 (GraphPad Software Inc.). Mathematica software (Wolfram Research Inc.) was used to calculate the Gaussian propagation of uncertainty. All quantitative data were tested for normal distribution using Shapiro-Wilk test. Parametric and non-parametric tests were used as appropriate. All statistical tests used in each experiment are listed in Supplementary Data S2. We used two-tailed Student’s t-test, two-tailed Mann-Whitney test, and paired two-tailed Student’s t-test. Differences were considered statistically significant at *P*-values below 0.05 and effect sizes (Cohen’s d for Student’s t-test, common language effect size θ for Mann-Whitney test) including the significance boundary (*d_c_* = 1, *θ_c_* = 0.5) within their uncertainty. \**P* < 0.05, \*\**P* < 0.01, \*\*\**P* < 0.001, \*\*\*\**P* < 0.0001. The *P*-value, effect size, and respective uncertainty for each experimental group is given in the figures or Supplementary Data S2. Respective effect sizes were determined according to (*77*) and (*78*) using custom scripts in Mathematica software (Wolfram Research Inc.). Unless indicated, variance for all groups data is presented as ± standard error of the mean (SEM). The sample size (n) for each experimental group is given in the figures or Supplementary Data S2. Graphs were generated using GraphPad Prism 9 or Mathematica software.

## SUPPLEMENTARY MATERIAL

Figs. S1 to S9

Tables S1 to S3

Data S1 to S2

## REFERENCES

1. S. Blackshaw, Why Has the Ability to Regenerate Following CNS Injury Been Repeatedly Lost Over the Course of Evolution? Front Neurosci 16, 831062 (2022).

2. D. O. Dias, C. Göritz, Fibrotic scarring following lesions to the central nervous system. Matrix biology : journal of the International Society for Matrix Biology 68-69, 561–570 (2018).

3. I. Francos-Quijorna et al., Chondroitin sulfate proteoglycans prevent immune cell phenotypic conversion and inflammation resolution via TLR4 in rodent models of spinal cord injury. Nature communications 13, 2933 (2022).

4. E. J. Bradbury, E. R. Burnside, Moving beyond the glial scar for spinal cord repair. Nature communications 10, 3879 (2019).

5. A. P. Tran, P. M. Warren, J. Silver, New insights into glial scar formation after spinal cord injury. Cell and tissue research, (2021).

6. E. J. Bradbury et al., Chondroitinase ABC promotes functional recovery after spinal cord injury. Nature 416, 636–640 (2002).

7. W. B. Cafferty, P. Duffy, E. Huebner, S. M. Strittmatter, MAG and OMgp synergize with Nogo-A to restrict axonal growth and neurological recovery after spinal cord trauma. The Journal of neuroscience : the official journal of the Society for Neuroscience 30, 6825–6837 (2010).

8. C. C. Stichel et al., Inhibition of collagen IV deposition promotes regeneration of injured CNS axons. The European journal of neuroscience 11, 632–646 (1999).

9. E. Moeendarbary et al., The soft mechanical signature of glial scars in the central nervous system. Nature communications 8, 14787 (2017).

10. X. Yu, R. V. Bellamkonda, Dorsal root ganglia neurite extension is inhibited by mechanical and chondroitin sulfate-rich interfaces. Journal of neuroscience research 66, 303–310 (2001).

11. H. J. Baumann et al., Softening of the chronic hemi-section spinal cord injury scar parallels dysregulation of cellular and extracellular matrix content. J Mech Behav Biomed Mater 110, 103953 (2020).

12. V. Tsata et al., A switch in pdgfrb+ cell-derived ECM composition prevents inhibitory scarring and promotes axon regeneration in the zebrafish spinal cord. Developmental cell 56, 509–524.e509 (2021).

13. D. Wehner et al., Wnt signaling controls pro-regenerative Collagen XII in functional spinal cord regeneration in zebrafish. Nature communications 8, 126 (2017).

14. M. H. Mokalled et al., Injury-induced ctgfa directs glial bridging and spinal cord regeneration in zebrafish. Science 354, 630–634 (2016).

15. C. G. Becker et al., L1.1 is involved in spinal cord regeneration in adult zebrafish. The Journal of neuroscience : the official journal of the Society for Neuroscience 24, 7837–7842 (2004).

16. V. Kroehne, D. Freudenreich, S. Hans, J. Kaslin, M. Brand, Regeneration of the adult zebrafish brain from neurogenic radial glia-type progenitors. Development 138, 4831–4841 (2011).

17. J. Ohnmacht et al., Spinal motor neurons are regenerated after mechanical lesion and genetic ablation in larval zebrafish. Development 143, 1464–1474 (2016).

18. P. Nauroy, S. Hughes, A. Naba, F. Ruggiero, The in-silico zebrafish matrisome: A new tool to study extracellular matrix gene and protein functions. Matrix biology : journal of the International Society for Matrix Biology, (2017).

19. Y. M. Yu et al., The extracellular matrix glycoprotein tenascin-C promotes locomotor recovery after spinal cord injury in adult zebrafish. Neuroscience 183, 238–250 (2011).

20. T. M. Tsarouchas et al., Dynamic control of proinflammatory cytokines Il-1beta and Tnf-alpha by macrophages in zebrafish spinal cord regeneration. Nature communications 9, 4670 (2018).

21. C. M. Nelson et al., Glucocorticoids Target Ependymal Glia and Inhibit Repair of the Injured Spinal Cord. Front Cell Dev Biol 7, 56 (2019).

22. J. Schweitzer et al., Tenascin-C is involved in motor axon outgrowth in the trunk of developing zebrafish. Developmental dynamics : an official publication of the American Association of Anatomists 234, 550–566 (2005).

23. Y. Li et al., Microglia-organized scar-free spinal cord repair in neonatal mice. Nature, (2020).

24. C. Y. Lin, Y. S. Lee, V. W. Lin, J. Silver, Fibronectin inhibits chronic pain development after spinal cord injury. Journal of neurotrauma 29, 589–599 (2012).

25. V. J. Tom, C. M. Doller, A. T. Malouf, J. Silver, Astrocyte-associated fibronectin is critical for axonal regeneration in adult white matter. The Journal of neuroscience : the official journal of the Society for Neuroscience 24, 9282–9290 (2004).

26. J. Chen et al., The extracellular matrix glycoprotein tenascin-C is beneficial for spinal cord regeneration. Mol Ther 18, 1769–1777 (2010).

27. K. S. O’Shea, L. H. Liu, V. M. Dixit, Thrombospondin and a 140 kd fragment promote adhesion and neurite outgrowth from embryonic central and peripheral neurons and from PC12 cells. Neuron 7, 231–237 (1991).

28. E. R. Bray et al., Thrombospondin-1 Mediates Axon Regeneration in Retinal Ganglion Cells. Neuron 103, 642–657 e647 (2019).

29. S. P. Hui, A. Dutta, S. Ghosh, Cellular response after crush injury in adult zebrafish spinal cord. Developmental dynamics : an official publication of the American Association of Anatomists 239, 2962–2979 (2010).

30. A. Didangelos et al., Rats and axolotls share a common molecular signature after spinal cord injury enriched in collagen-1. bioRxiv, (2017).

31. N. Klapka, H. W. Muller, Collagen matrix in spinal cord injury. Journal of neurotrauma 23, 422–435 (2006).

32. R. Kalluri, Basement membranes: structure, assembly and role in tumour angiogenesis. Nat Rev Cancer 3, 422–433 (2003).

33. A. Didangelos et al., High-throughput proteomics reveal alarmins as amplifiers of tissue pathology and inflammation after spinal cord injury. Scientific reports 6, 21607 (2016).

34. J. Tica, E. J. Bradbury, A. Didangelos, Combined Transcriptomics, Proteomics and Bioinformatics Identify Drug Targets in Spinal Cord Injury. International journal of molecular sciences 19, (2018).

35. X. Pang, N. Dong, Z. Zheng, Small Leucine-Rich Proteoglycans in Skin Wound Healing. Front Pharmacol 10, 1649 (2020).

36. R. Kuranov et al., Complementary use of cross-polarization and standard OCT for differential diagnosis of pathological tissues. Optics express 10, 707–713 (2002).

37. J. M. Schmitt, S. H. Xiang, Cross-polarized backscatter in optical coherence tomography of biological tissue. Optics letters 23, 1060–1062 (1998).

38. R. Prevedel, A. Diz-Munoz, G. Ruocco, G. Antonacci, Brillouin microscopy: an emerging tool for mechanobiology. Nature methods 16, 969–977 (2019).

39. G. Antonacci et al., Recent progress and current opinions in Brillouin microscopy for life science applications. Biophys Rev 12, 615–624 (2020).

40. G. Scarcelli et al., Noncontact three-dimensional mapping of intracellular hydromechanical properties by Brillouin microscopy. Nature methods 12, 1132–1134 (2015).

41. C. Riquelme-Guzman et al., In vivo assessment of mechanical properties during axolotl development and regeneration using confocal Brillouin microscopy. Open Biol 12, 220078 (2022).

42. C. Bevilacqua, H. Sanchez-Iranzo, D. Richter, A. Diz-Munoz, R. Prevedel, Imaging mechanical properties of sub-micron ECM in live zebrafish using Brillouin microscopy. Biomed Opt Express 10, 1420–1431 (2019).

43. R. Schlüssler et al., Mechanical Mapping of Spinal Cord Growth and Repair in Living Zebrafish Larvae by Brillouin Imaging. Biophysical journal 115, 911–923 (2018).

44. S. Möllmert et al., Zebrafish Spinal Cord Repair Is Accompanied by Transient Tissue Stiffening. Biophysical journal 118, 448–463 (2020).

45. K. R. Long et al., Extracellular Matrix Components HAPLN1, Lumican, and Collagen I Cause Hyaluronic Acid-Dependent Folding of the Developing Human Neocortex. Neuron 99, 702–719 e706 (2018).

46. M. Brand, Granato, M. and Nüsslein-Volhard, C, C. N.-V. a. R. Dahm, Ed. (Oxford University Press, Oxford, 2002).

47. P. Alestrom et al., Zebrafish: Housing and husbandry recommendations. Lab Anim, 23677219869037 (2019).

48. K. Ando et al., Clarification of mural cell coverage of vascular endothelial cells by live imaging of zebrafish. Development 143, 1328–1339 (2016).

49. K. Asakawa et al., Genetic dissection of neural circuits by Tol2 transposon-mediated Gal4 gene and enhancer trapping in zebrafish. Proceedings of the National Academy of Sciences of the United States of America 105, 1255–1260 (2008).

50. J. M. Davison et al., Transactivation from Gal4-VP16 transgenic insertions for tissuespecific cell labeling and ablation in zebrafish. Developmental biology 304, 811–824 (2007).

51. J. Freeman et al., Mapping brain activity at scale with cluster computing. Nature methods 11, 941–950 (2014).

52. D. Wehner, C. Jahn, G. Weidinger, Use of the TetON System to Study Molecular Mechanisms of Zebrafish Regeneration. Journal of visualized experiments : JoVE, e52756 (2015).

53. F. Knopf et al., Dually inducible TetON systems for tissue-specific conditional gene expression in zebrafish. Proceedings of the National Academy of Sciences of the United States of America 107, 19933–19938 (2010).

54. M. L. Suster, H. Kikuta, A. Urasaki, K. Asakawa, K. Kawakami, Transgenesis in zebrafish with the tol2 transposon system. Methods in molecular biology 561, 41–63 (2009).

55. N. John, J. Kolb, D. Wehner, Mechanical spinal cord transection in larval zebrafish and subsequent whole-mount histological processing. STAR Protocols 3, 101093 (2022).

56. H. B. Schiller et al., Time-and compartment-resolved proteome profiling of the extracellular niche in lung injury and repair. Mol Syst Biol 11, 819 (2015).

57. J. Cox, M. Mann, MaxQuant enables high peptide identification rates, individualized p.p.b.-range mass accuracies and proteome-wide protein quantification. Nat Biotechnol 26, 1367–1372 (2008).

58. S. Tyanova, J. Cox, Perseus: A Bioinformatics Platform for Integrative Analysis of Proteomics Data in Cancer Research. Methods in molecular biology 1711, 133–148 (2018).

59. V. G. Tusher, R. Tibshirani, G. Chu, Significance analysis of microarrays applied to the ionizing radiation response. Proceedings of the National Academy of Sciences of the United States of America 98, 5116–5121 (2001).

60. U. Raudvere et al., g:Profiler: a web server for functional enrichment analysis and conversions of gene lists (2019 update). Nucleic Acids Res 47, W191–W198 (2019).

61. B. Jassal et al., The reactome pathway knowledgebase. Nucleic Acids Res 48, D498–D503 (2020).

62. A. Didangelos, B. Roschitzki. (Mendeley Data, 2018).

63. J. C. Oliveros. (2007-2015).

64. J. Jablonski et al., Experimental Epileptogenesis in a Cell Culture Model of Primary Neurons from Rat Brain: A Temporal Multi-Scale Study. Cells 10, (2021).

65. G. R. Hartl, A. Parmar, G. Sharma, K. Singh, Cross-Polarized Optical Coherence Tomography System with Unpolarized Light. Photonics 9, 76 (2022).

66. K. Singh, G. Sharma, G. J. Tearney, Estimation and compensation of dispersion for a high-resolution optical coherence tomography system. Journal of Optics 20, 025301 (2018).

67. G. M. R. De Luca et al., Re-scan confocal microscopy: scanning twice for better resolution. Biomed. Opt. Express 4, 2644–2656 (2013).

68. T. Boothe et al., A tunable refractive index matching medium for live imaging cells, tissues and model organisms. eLife 6, (2017).

69. K. Kim, J. Guck, The Relative Densities of Cytoplasm and Nuclear Compartments Are Robust against Strong Perturbation. Biophysical journal 119, 1946–1957 (2020).

70. E. Cuche, P. Marquet, C. Depeursinge, Spatial filtering for zero-order and twin-image elimination in digital off-axis holography. Applied optics 39, 4070–4075 (2000).

71. E. Wolf, Three-dimensional structure determination of semi-transparent objects from holographic data. Optics Communications 1, 153–156 (1969).

72. Y. Sung et al., Optical diffraction tomography for high resolution live cell imaging. Optics express 17, 266–277 (2009).

73. K. Kim et al., High-resolution three-dimensional imaging of red blood cells parasitized by Plasmodium falciparum and in situ hemozoin crystals using optical diffraction tomography. J Biomed Opt 19, 011005 (2014).

74. C. Arshadi, U. Gunther, M. Eddison, K. I. S. Harrington, T. A. Ferreira, SNT: a unifying toolbox for quantification of neuronal anatomy. Nature methods 18, 374–377 (2021).

75. R. Barer, Interference Microscopy and Mass Determination. 169, 366–367 (1952).

76. H. Zhao, P. H. Brown, P. Schuck, On the distribution of protein refractive index increments. Biophysical journal 100, 2309–2317 (2011).

77. M. P. Perme, D. Manevski, Confidence intervals for the Mann–Whitney test. Statistical Methods in Medical Research 28, 3755–3768 (2019).

78. J. Cohen. J. Cohen, Ed., Statistical Power Analysis for the Behavioral Sciences (Routledge, New York, 1988).

