## Supplementary information for "Small leucine-rich proteoglycans inhibit CNS regeneration by modifying the structural and mechanical properties of the lesion environment"

#### TITLE

#### AFFILIATIONS

### Supplementary Information

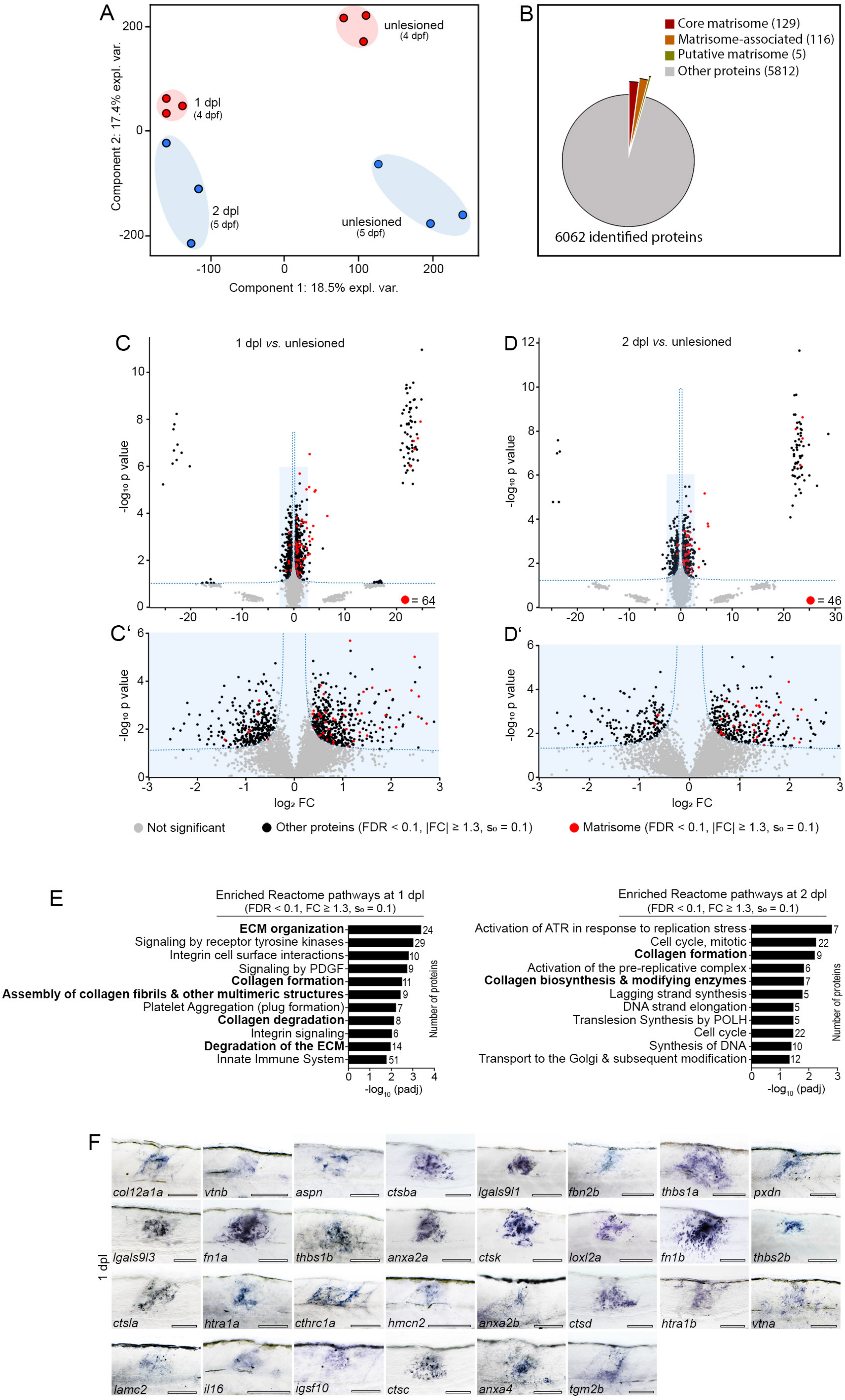
A
unlesioned (4 dpf)
1 dpl (4 dpf)
2 dpl (5 dpf)
unlesioned (5 dpf)
Component 2: 17.4% expl. var.
Component 1: 18.5% expl. var.
B
Core matrisome (129)
Matrisome-associated (116)
Putative matrisome (5)
Other proteins (5812)
6062 identified proteins
C
1 dpl vs. unlesioned
= 64
D
2 dpl vs. unlesioned
= 46
C'
D'
-log10 p value
log2 FC
Not significant
Other proteins (FDR < 0.1, |FC| ≥ 1.3, s0 = 0.1)
Matrisome (FDR < 0.1, |FC| ≥ 1.3, s0 = 0.1)
E
Enriched Reactome pathways at 1 dpl
(FDR < 0.1, FC ≥ 1.3, s0 = 0.1)
ECM organization 24
Signaling by receptor tyrosine kinases 29
Integrin cell surface interactions 10
Signaling by PDGF 9
Collagen formation 11
Assembly of collagen fibrils & other multimeric structures 9
Platelet Aggregation (plug formation) 7
Collagen degradation 8
Integrin signaling 6
Degradation of the ECM 14
Innate Immune System 51
Number of proteins
-log10 (padj)
Enriched Reactome pathways at 2 dpl
(FDR < 0.1, FC ≥ 1.3, s0 = 0.1)
Activation of ATR in response to replication stress 7
Cell cycle, mitotic 22
Collagen formation 9
Activation of the pre-replicative complex 6
Collagen biosynthesis & modifying enzymes 7
Lagging strand synthesis 5
DNA strand elongation 5
Translesion Synthesis by POLH 5
Cell cycle 22
Synthesis of DNA 10
Transport to the Golgi & subsequent modification 12
Number of proteins
-log10 (padj)
F
1 dpl
col12a1a
vtnb
aspn
ctsba
lgals9l1
fbn2b
thbs1a
pxdn
lgals9l3
fn1a
thbs1b
anxa2a
ctsk
loxl2a
fn1b
thbs2b
ctsla
htra1a
cthrc1a
hmcn2
anxa2b
ctsd
htra1b
vtna
lamc2
il16
igsf10
ctsc
anxa4
tgm2b

**Figure S1 | Mass spectrometry-based quantitative proteomics reveals changes in ECM composition during zebrafish spinal cord regeneration.**

**A)** Principle component analysis of mass spectrometry-based proteomics data using quantitative values of all identified proteins across samples. Lesioned samples (1 dpl, 2 dpl) cluster distinct from unlesioned age-matched control samples (4 dpf, 5 dpf). Each data point represents one biological replicate.

**B)** Pie chart showing the total number of identified proteins grouped by core matrisome proteins, matrisome-associated proteins, putative matrisome proteins, and other proteins.

**C-D)** Volcano plots of all quantified proteins for the given analyses (1 dpl *vs*. unlesioned age-matched controls, C; 2 dpl *vs.* unlesioned age-matched controls, D) with their $log_2$-transformed ratios of the mean-centered abundances (FC, fold change) and $-log_{10}$-transformed *P*-values (two-sided t-test). Dashed line indicates the threshold of a permutation-based FDR correction for multiple hypotheses ($FDR < 0.1$, $s_0 = 0.1$) for identification of significantly altered abundances. Proteins with significantly altered abundance were further filtered by $|FC| \geq 1.3$. Area highlighted (blue) in (C) and (D) is shown at a different scale in (C') and (D').

**E)** Reactome pathway analysis of differentially enriched proteins reveals ECM-associated terms (in bold) being overrepresented at 1 dpl and 2 dpl.

**F)** Expression of indicated genes coding for differentially regulated matrisome proteins is upregulated in the lesion site at 1 dpl, as determined by *in situ* hybridization (lateral view; rostral is left). $n \geq 9$ for each gene. Scale bars: 100 µm.

**A-F)** dpf, days post-fertilization; dpl, days post-lesion; expl. var., explained variance; FC, fold change; FDR, false discovery rate.

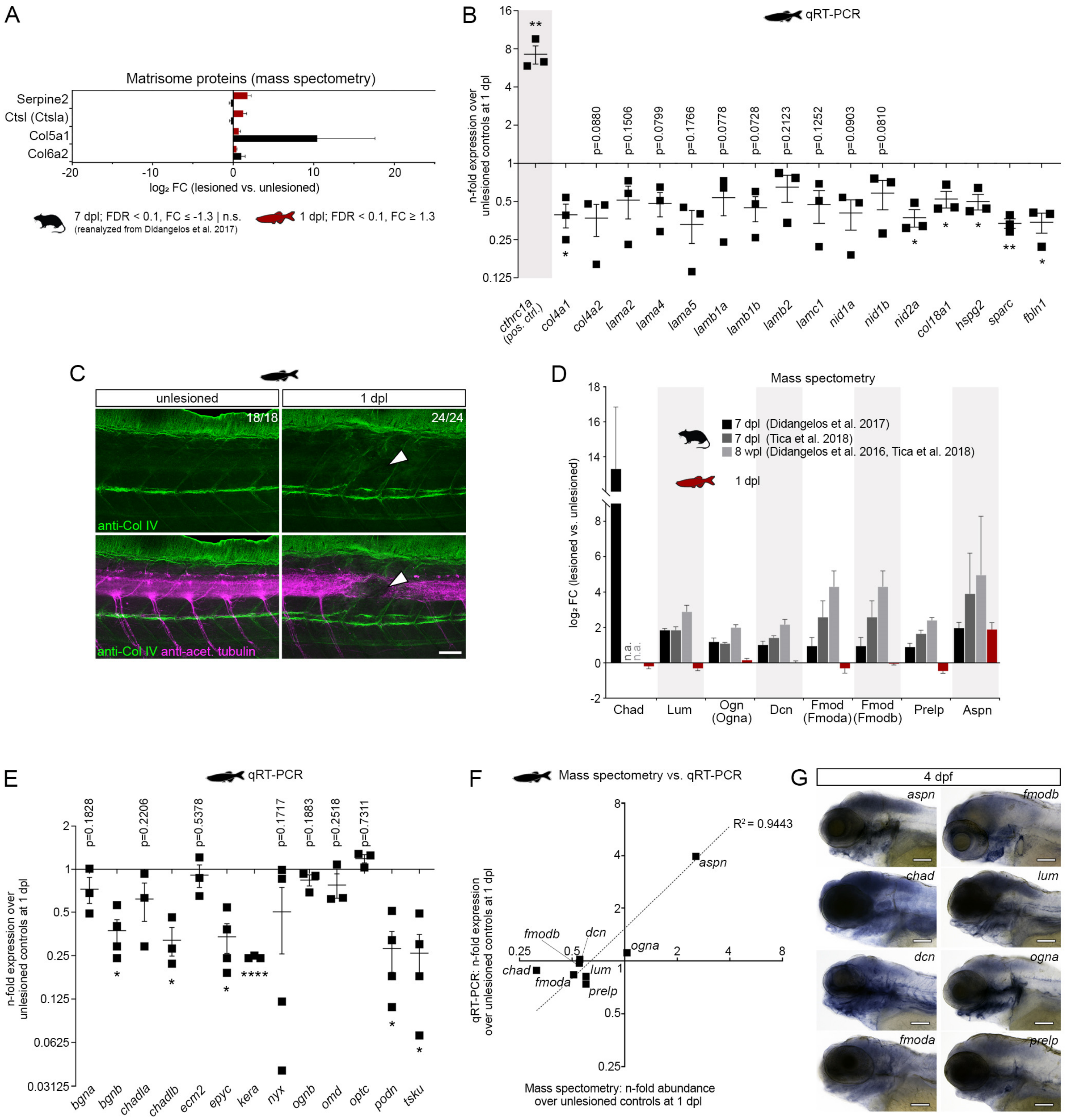

**Figure S2 | Basal lamina constituents and small leucine-rich proteoglycans are not enriched in the zebrafish spinal lesion site.**

**A)** Comparative proteomics analysis reveals differentially enriched matrisome proteins between rat (black) and zebrafish (red) after SCI. Shown are proteins that exhibit a high abundance (FDR < 0.1, FC ≥ 1.3) after SCI in zebrafish but a low abundance (FDR < 0.1, FC ≤ 1.3 | n.s.) in the rat spinal lesion site. Data are means ± SEM.

**B)** Fold change expression of indicated genes in the zebrafish spinal lesion site at 1 dpl over unlesioned age-matched controls, as determined by qRT-PCR. Expression of indicated genes coding for basal lamina constituents is not upregulated in the zebrafish spinal lesion site at 1 dpl.

Note that *cthrc1a* is not a component of the basal lamina complex and served as a positive control for the detection of injury-induced gene expression (*1*). Fold change values are presented in log scale. Each data point represents one biological replicate. Data are means ± SEM.

**C)** Immunolabeling of type IV collagen is not increased in the zebrafish spinal lesion site at 1 dpl (arrowheads). Spinal axons are labeled with immunofluorescence against acetylated tubulin (magenta). The number of specimens displaying the phenotype and the total number of experimental specimens is given. Images shown are maximum intensity projections of unlesioned trunk or lesion site (lateral view; rostral is left).

**D)** Comparative analysis of indicated proteomics datasets reveals differential enrichment of Chad, Dcn, Fmoda, Fmodb, Lum, Ogna, and Prelp proteins after SCI in rat (grey tones) and zebrafish (red). Note that Aspn is enriched after SCI both in rat and zebrafish. Data are means ± SEM.

**E)** Fold change expression of indicated genes in the zebrafish spinal lesion site at 1 dpl over unlesioned age-matched controls, as determined by qRT-PCR. Expression of genes coding for indicated SLRP proteins is not upregulated at 1 dpl. Fold change values are presented in log scale. Each data point represents one biological replicate. Data are means ± SEM.

**F)** Mean fold change values of indicated genes and proteins at 1 dpl over unlesioned age-matched controls as determined by qRT-PCR and mass spectrometry-based quantitative proteomics, are highly correlated ($R^2 = 0.9443$). Fold change values are presented in log scale.

**G)** Transcripts of indicated genes (blue) are detectable by *in situ* hybridization in whole-mount zebrafish larvae at 4 dpf. Images shown are lateral views of the heads of animals depicted in Fig. 2C.

**A-G)** $^*P < 0.05$, $^{**}P < 0.01$, $^{****}P < 0.0001$. Scale bars: 100 µm (G), 50 µm (C). dpf, days post-fertilization; dpl, days post-lesion; FC, fold change; FDR, false discovery rate; n.s., not significant.

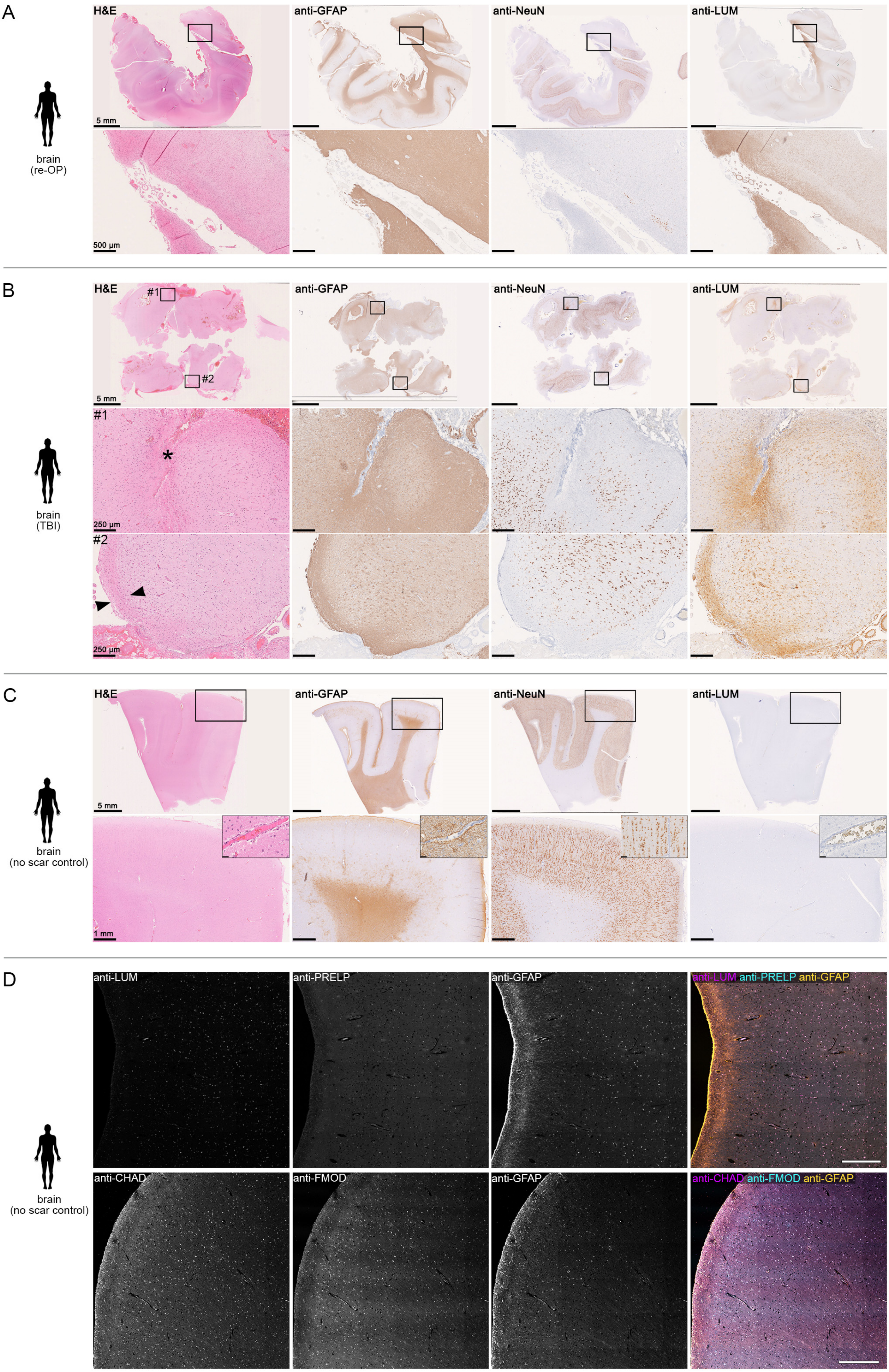
A
H&E
anti-GFAP
anti-NeuN
anti-LUM
5 mm
500 µm
brain
(re-OP)
B
#1
#2
250 µm
brain
(TBI)
C
1 mm
brain
(no scar control)
D
anti-PRELP
anti-CHAD
anti-FMOD
anti-LUM anti-PRELP anti-GFAP
anti-CHAD anti-FMOD anti-GFAP

**Figure S3 | SLRPs are enriched in human brain lesions.**

**A-C)** Immunohistochemical examination of human brain specimens. Hematoxylin & eosin (H&E) staining, and 3,3′-Diaminobenzidine (DAB) staining of anti-GFAP, anti-NeuN, and anti-LUM antibodies on scarred brain tissue from patients with previous surgery (re-OP; A) or traumatic brain injury (TBI; B), and no scar control brain tissue (C). Areas of scarring were identified by H&E staining pattern and absence of immunoreactivity of the neuronal marker anti-NeuN. Anti-LUM immunoreactivity is increased in areas of scarring caused by previous surgery (A), contusion (arrowheads in B), or local hemorrhage (asterisk in B). Anti-LUM immunoreactivity is negligible in healthy human brain autopsy tissue with no signs of scarring (C). Shown are coronal sections. Size of scale bars is given in the figure.

**D)** Anti-LUM, anti-PRELP, anti-CHAD and anti-FMOD immunoreactivity is negligible in healthy human brain autopsy tissue with no signs of scarring. Images shown are immunofluorescence controls for data shown in Fig. 3. Shown are coronal sections. Scale bars: 500 µm.

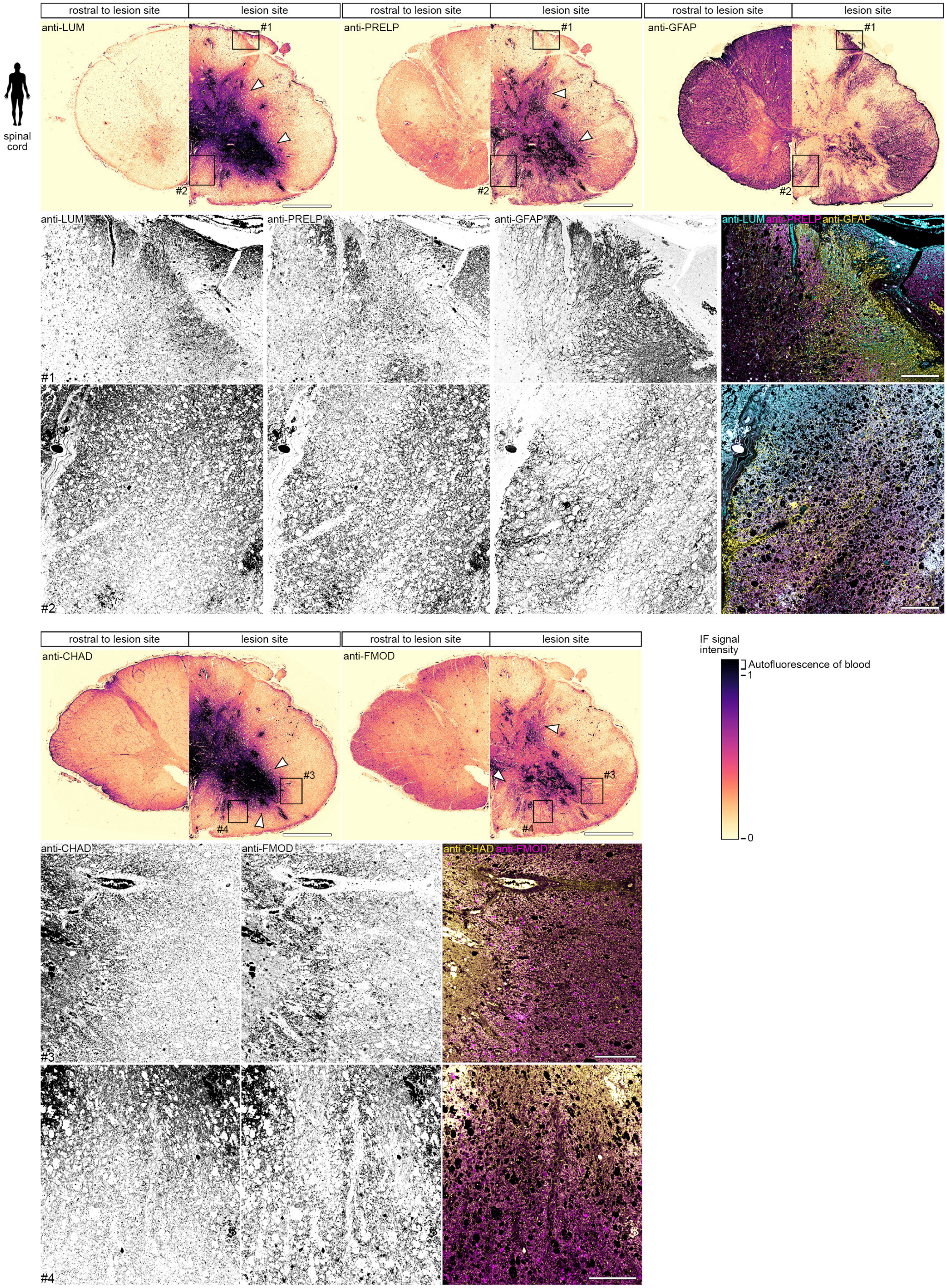
rostral to lesion site
lesion site
anti-LUM
anti-PRELP
anti-GFAP
spinal cord
#1
#2
anti-LUM anti-PRELP anti-GFAP
#3
#4
anti-CHAD
anti-FMOD
anti-CHAD anti-FMOD
IF signal intensity
Autofluorescence of blood
1
0

**Figure S4 | SLRPs are enriched in human spinal cord lesions.**

Anti-LUM, anti-PRELP, anti-CHAD, and anti-FMOD immunoreactivity is increased (arrowheads; see calibration bar of lookup table) in the epicenter of the injured human spinal cord at nine days-post injury, as compared to rostral control segments of the same patient. Note that immunoreactivity is mainly observed in proximity to the hemorrhage (black; see calibration bar of LUT). Also note that with the exception of blood-derived autofluorescence, individual channels show distinct pattern of immunofluorescence (IF) signal (insets). Shown are transversal sections (dorsal is up). Scale bars: 2 mm, 200 µm (insets).

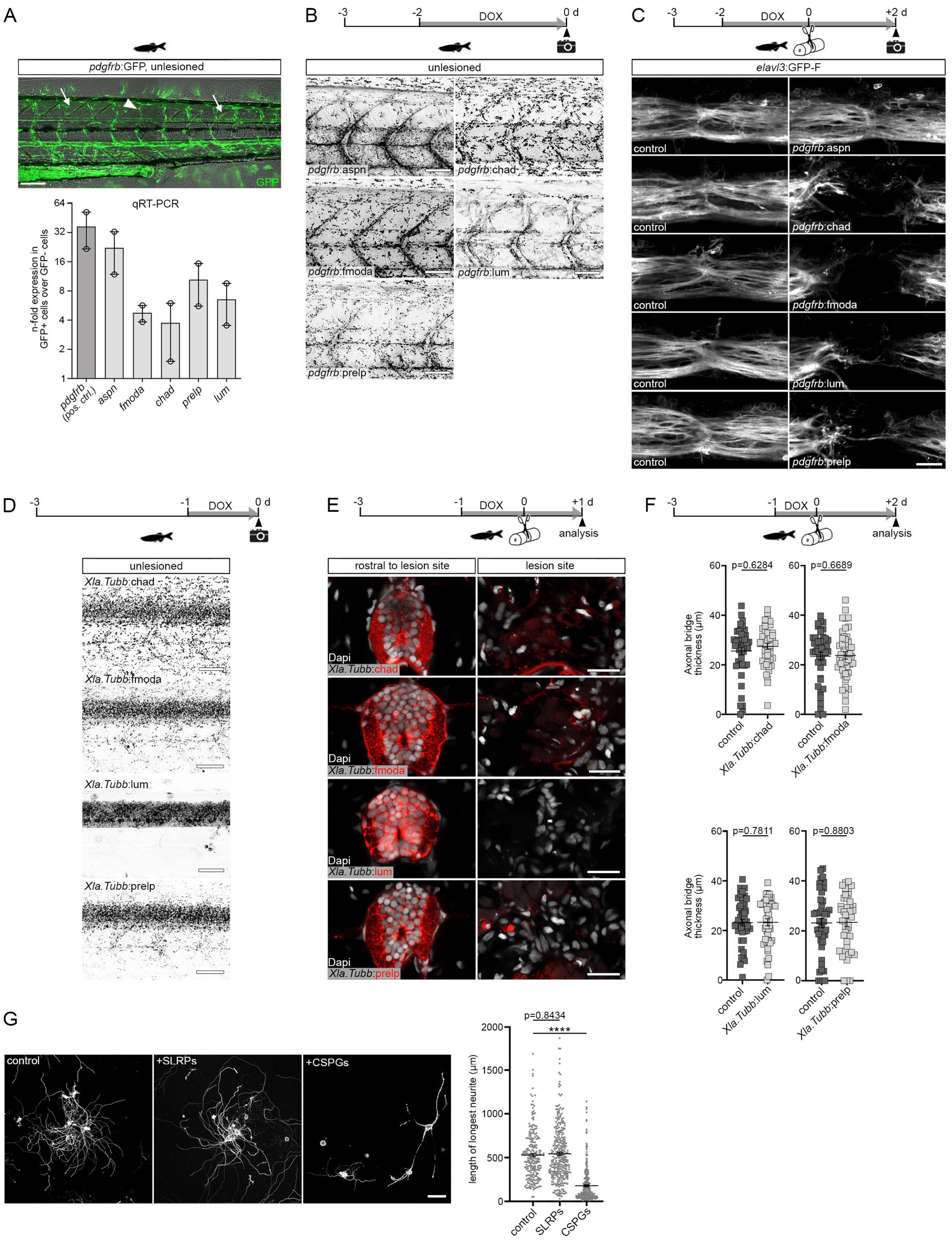
A
pdgfrb:GFP, unlesioned
GFP
qRT-PCR
n-fold expression in GFP+ cells over GFP- cells
pdgfrb (pos. ctrl.)
aspn
fmoda
chad
prelp
lum
B
DOX
0 d
unlesioned
pdgfrb:aspn
pdgfrb:chad
pdgfrb:fmoda
pdgfrb:lum
pdgfrb:prelp
C
DOX
+2 d
elavl3:GFP-F
control
pdgfrb:aspn
pdgfrb:chad
pdgfrb:fmoda
pdgfrb:lum
pdgfrb:prelp
D
DOX
0 d
unlesioned
Xla.Tubb:chad
Xla.Tubb:fmoda
Xla.Tubb:lum
Xla.Tubb:prelp
E
DOX
+1 d
analysis
rostral to lesion site
lesion site
Dapi
Xla.Tubb:chad
Xla.Tubb:fmoda
Xla.Tubb:lum
Xla.Tubb:prelp
F
DOX
+2 d
analysis
Axonal bridge thickness (µm)
p=0.6284
p=0.6689
p=0.7811
p=0.8803
control
Xla.Tubb:chad
Xla.Tubb:fmoda
Xla.Tubb:lum
Xla.Tubb:prelp
G
control
+SLRPs
+CSPGs
length of longest neurite (µm)
p=0.8434
****
control
SLRPs
CSPGs

**Figure S5 | Cell type-specific targeting of SLRPs in zebrafish, and SLRPs do not act on neurons to inhibit neurite extension.**

**A)** *pdgfrb*$^+$ myoseptal (arrows) and perivascular (arrowhead) cells (green) are a major source of endogenous SLRPs in larval zebrafish. Shown is the fold change expression of indicated genes in FACS-isolated GFP$^+$ cells over GFP$^-$ cells in trunk tissue of *pdgfrb*:GFP transgenic animals at 4 dpf, as determined by qRT-PCR. Note that *pdgfrb* served as control for enrichment of *pdgfrb*$^+$ cells. Fold change values are presented in log scale. Each data point represents one biological replicate.

**B)** mCherry fluorescence (black) is selectively induced in *pdgfrb*$^+$ cells of indicated *pdgfrb*:TetA; *TetRE*:SLRP-mCherry (short *pdgfrb*:SLRP) transgenic zebrafish following DOX treatment. Note that the SLRP-mCherry fusions are secreted proteins and that the pattern of mCherry fluorescence resembles that of GFP in *pdgfrb*:GFP transgenic animals (see A). Also note that the pattern of protein deposition varies among different SLRPs and that some exhibit fiber-like structures.

**C)** *pdgfrb*$^+$ cell-specific induction of the SLRPs *chad*, *fmoda*, *lum*, and *prelp* but not *aspn* in *pdgfrb*:SLRP transgenic zebrafish reduces the thickness of the axonal bridge (white; analyzed in *elavl3*:GFP-F transgenics). Shown are example images of axonal bridges quantified in Fig. 4B'.

**D)** mCherry fluorescence (black) is selectively induced in neurons of indicated *Xla.Tubb*:TetA; *TetRE*:SLRP-mCherry (short *Xla.Tubb*:SLRP) transgenic zebrafish following DOX treatment. Note that the SLRP-mCherry fusions are secreted proteins. Also note that the mCherry fluorescence is primarily confined to the spinal cord.

**E)** Neuron-specific induction of indicated *slrp-mCherry* fusions in *Xla.Tubb*:TetA;*TetRE*:SLRP-mCherry (short *Xla.Tubb*:SLRP) transgenic zebrafish leads to negligible mCherry fluorescence (red) in the spinal lesion site at 1 dpl. Note that strong mCherry fluorescence (red) is detectable in intact spinal cord tissue rostral to the lesion site.

**F)** Neuron-specific induction of *chad*, *fmoda*, *lum,* or *prelp* in *pdgfrb*:SLRP transgenic zebrafish does not reduce the thickness of the axonal bridge (analyzed in *elavl3*:GFP-F transgenics) at 2 dpl. Each data point represents one animal.

**G)** Neurite outgrowth is not reduced when primary adult murine dorsal root ganglion neurons are cultured on substrates coated with a mixture of human CHAD, FMOD, LUM, and PRELP proteins, as compared to controls. Neurite growth is significantly reduced in the presence of CSPGs.

**A-G)** Images shown are maximum intensity projections of unlesioned trunk or lesion site (A-D; lateral view; rostral is left), or transverse views of unlesioned trunk or lesion site (E; dorsal is up). Data are means ± SEM. ****$P < 0.0001$. Scale bars: 100 µm (A, G), 50 µm (B, D), 25 µm (C), 20 µm (E). d, days; DOX, doxycycline.

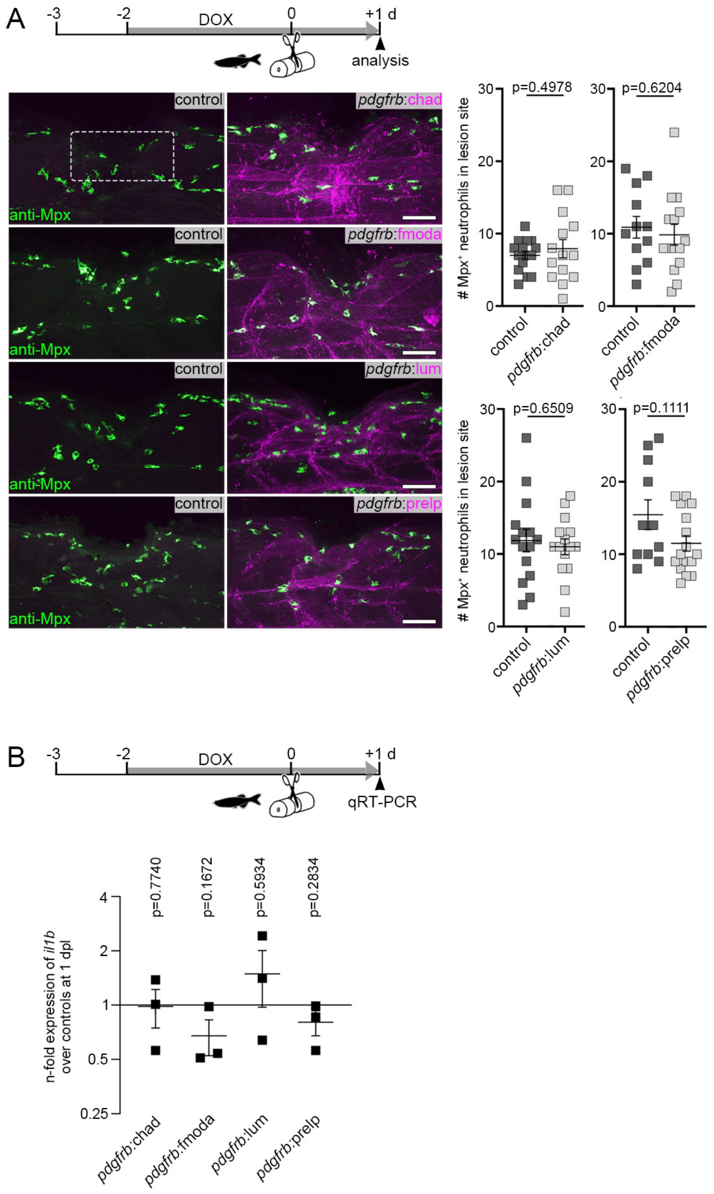


**Figure S6 | Targeting SLRPs to the injury ECM does not prevent inflammation resolution.**

**A)** Induction of *chad*, *fmoda*, *lum,* or *prelp* in *pdgfrb*:SLRP transgenic zebrafish does not lead to a higher number of Mpx$^+$ neutrophils in the lesion site at 1 dpl. Images shown are maximum intensity projections of the lesion site (lateral view; rostral is left). The dashed rectangle indicates the region of quantification.

**B)** Fold change expression of *il1b* in the spinal lesion site of *pdgfrb*:SLRP transgenic zebrafish over controls at 1 dpl, as determined by qRT-PCR. Induction of *chad*, *fmoda*, *lum,* or *prelp* in *pdgfrb*:SLRP transgenics does not lead to increased expression levels of the proinflammatory cytokine *il1b*. Fold change values are presented in log scale.

**A-B)** Each data point represents one animal. Data are means ± SEM. Scale bars: 50 µm. d, days; dpl, days post-lesion; DOX, doxycycline.

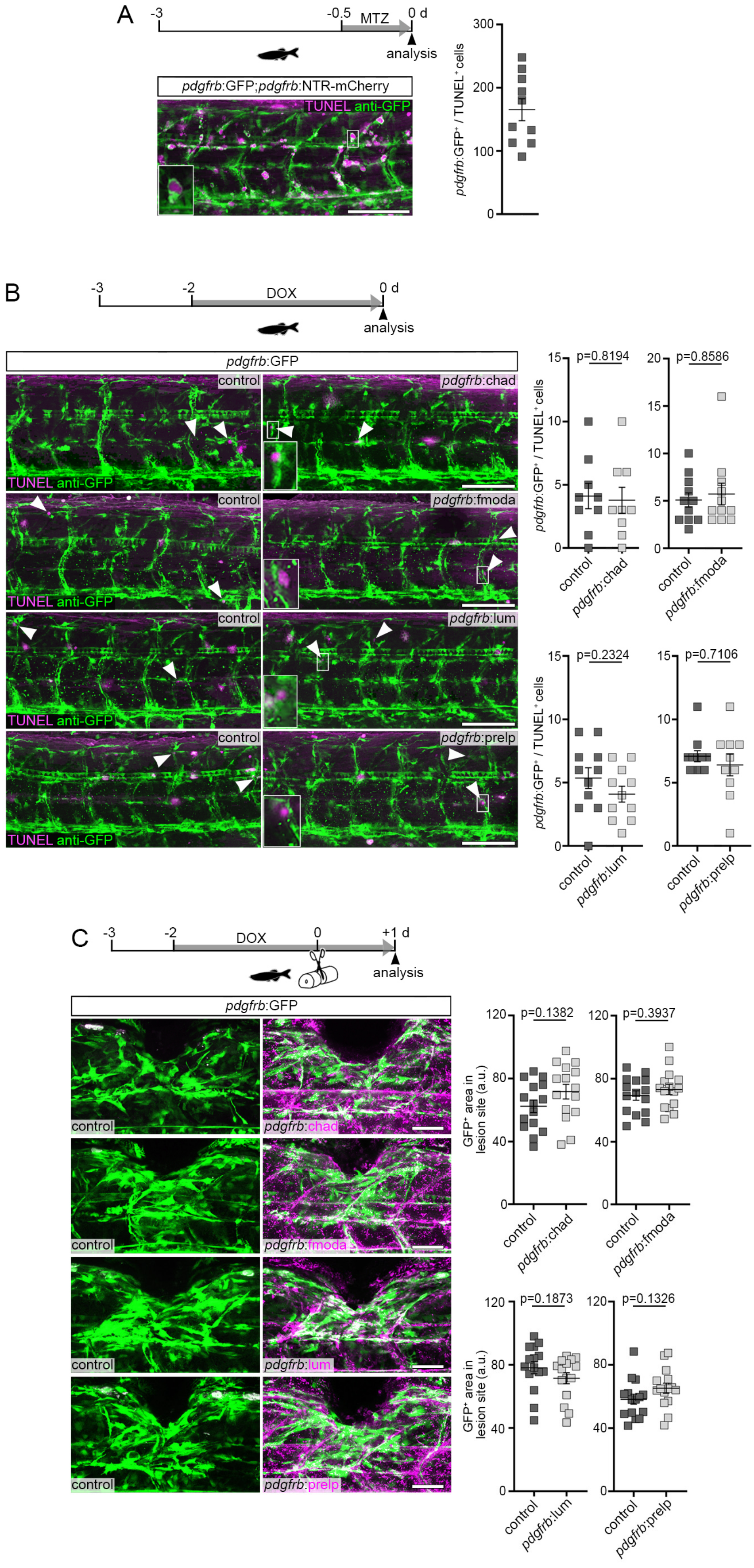


**Figure S7 | *pdgfrb*⁺ cell-specific induction of SLRPs does not alter the fibroblast response.**

**A)** Detection of a high number of *pdgfrb*:GFP⁺/TUNEL⁺ (green/magenta) cells in *pdgfrb*:GFP;*pdgfrb*:NTR-mCherry transgenic zebrafish following metronidazole (MTZ) treatment, reveals efficient TUNEL probe penetration in whole-mount preparations.

**B)** Induction of *chad*, *fmoda*, *lum,* or *prelp* in *pdgfrb*:SLRP transgenic zebrafish does not increase the number of *pdgfrb*:GFP$^+$ cells (green) that undergo apoptosis (TUNEL$^+$; magenta).

**C)** Induction of *chad*, *fmoda*, *lum,* or *prelp* in *pdgfrb*:SLRP transgenic zebrafish does not prevent the appearance of *pdgfrb*$^+$ fibroblast-like cells (green) in the lesion site at 1 dpl.

**A-C)** Images shown are maximum intensity projections of the lesion site or unlesioned trunk (lateral view; rostral is left). Each data point represents one animal. Data are means ± SEM. Scale bars: 100 µm (A, B), 50 µm (C). a.u., arbitrary unit; d, days; DOX, doxycycline.

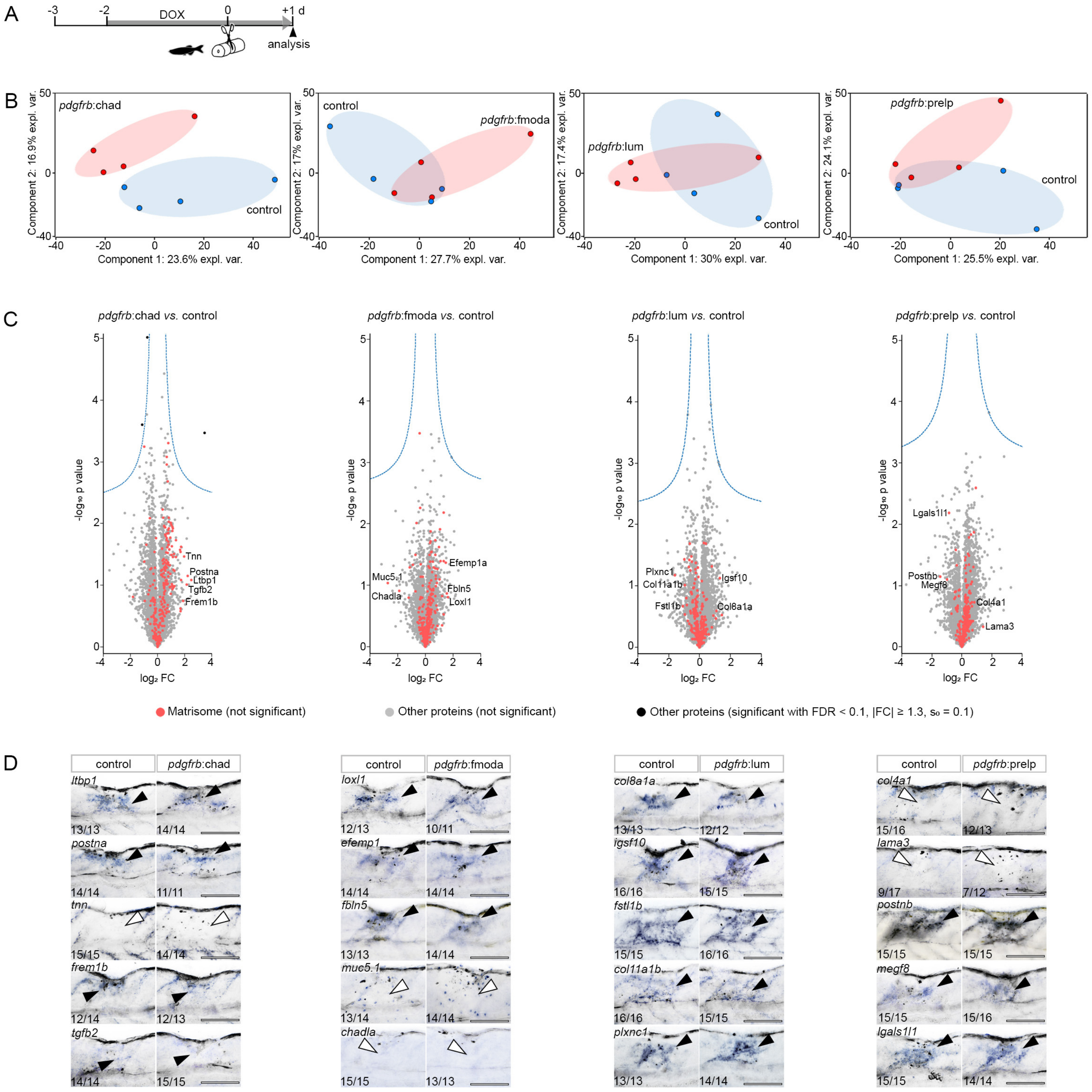


**Figure S8 | Targeting SLRPs to the injury ECM does not alter its composition.**

**A)** Timeline for experimental treatments shown in (B), (C), and (D). d, days.

**B)** Principle component analysis of mass spectrometry-based (MS) quantitative proteomics data from indicated experimental conditions, using quantitative values of all identified proteins across samples. Each data point represents one biological replicate. expl. var., explained variance.

**C)** Induction of *chad*, *fmoda*, *lum,* or *prelp* in *pdgfrb*:SLRP transgenic zebrafish does not lead to significant differences in the abundance of matrisome proteins at 1 dpl (red data points), as compared to controls. Shown are volcano plots of all quantified proteins for the given analyses with their $log_2$-transformed ratios of the mean-centered abundances (FC, fold change) and -$log_{10}$-transformed *P*-values (two-sided t-test). Dashed line indicates the threshold of a permutation-based FDR correction for multiple hypotheses (FDR < 0.1, $s_0 = 0.1$) for identification of significantly altered abundances. Proteins with significantly altered abundance were further filtered by $|FC| \geq 1.3$. Since the MS analysis cannot discriminate between endogenous and ectopic SLRP proteins, the respective manipulated protein is not shown in the volcano plot. Indicated proteins were selected for verification by *in situ* hybridization (ISH; see D).

**D)** Induction of *chad*, *fmoda*, *lum*, or *prelp* in *pdgfrb*:SLRP transgenic zebrafish does not lead to major changes in the expression (blue) of indicated genes in the lesion site at 1 dpl, as determined by ISH. Transcript levels of five genes coding for matrisome proteins that showed the most substantial evidence of regulation in each experimental condition in (C) were evaluated. The number of specimens displaying the phenotype and the total number of experimental specimens is given. Black arrowheads indicate ISH signal in the center of the lesion site, white arrowheads indicate absence of ISH signal. Images shown are brightfield recordings of the lesion site (lateral view; rostral is left). Scale bars: 100 μm.

A

$$M' = \frac{\lambda^2 \nu_{\mathrm{B}}^2 \rho}{4n^2} \quad (1)$$

$$\rho \approx \frac{n - n_{\mathrm{fluid}}}{\alpha} + \rho_{\mathrm{fluid}} \left( 1 - \frac{\theta (n - n_{\mathrm{fluid}})}{\alpha} \right) \quad (2)$$

B

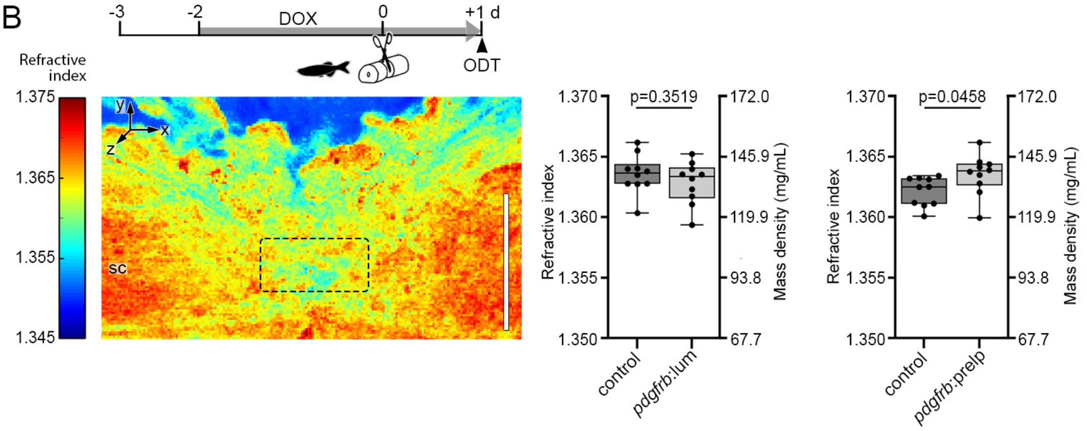


C

$$\Delta M' = \sqrt{\left( \frac{\partial M'}{\partial n} \Delta n \right)^2 + \left( \frac{\partial M'}{\partial \nu_{\mathrm{B}}} \Delta \nu_{\mathrm{B}} \right)^2 + \left( \frac{\partial M'}{\partial \alpha} \Delta \alpha \right)^2 + \left( \frac{\partial M'}{\partial \theta} \Delta \theta \right)^2} \quad (1)$$

$$\Delta M' = \frac{\lambda^2 \nu_{\mathrm{B}}}{4n^3 \alpha^2} \sqrt{(\xi_n \Delta n)^2 + (\xi_{\nu_{\mathrm{B}}} \Delta \nu_{\mathrm{B}})^2 + (\xi_\alpha \Delta \alpha)^2 + (\xi_\theta \Delta \theta)^2} \quad (2)$$

with...

$$\xi_n \equiv \alpha \nu_{\mathrm{B}} \left( n - 2n_{\mathrm{fluid}} + \rho_{\mathrm{fluid}} \left( 2\alpha - n\theta + 2n_{\mathrm{fluid}} \theta \right) \right) \quad (3)$$

$$\xi_{\nu_{\mathrm{B}}} \equiv 2n\alpha \left( n + \alpha \rho_{\mathrm{fluid}} - n\theta \rho_{\mathrm{fluid}} + n_{\mathrm{fluid}} \left( \theta \rho_{\mathrm{fluid}} - 1 \right) \right) \quad (4)$$

$$\xi_\alpha \equiv n \nu_{\mathrm{B}}^2 (\theta \rho_{\mathrm{fluid}} - 1)(n - n_{\mathrm{fluid}}) \quad (5)$$

$$\xi_\theta \equiv n \alpha \nu_{\mathrm{B}} \rho_{\mathrm{fluid}} (n - n_{\mathrm{fluid}}) \quad (6)$$

D

$$\delta_{\nu_{\mathrm{B}}} \equiv \frac{M'(\gamma \nu_{\mathrm{B}}, n, \theta)}{M'(\nu_{\mathrm{B}}, n, \theta)} = \gamma^2 \quad (1)$$

$$\delta_n \equiv \frac{M'(\nu_{\mathrm{B}}, \gamma n, \theta)}{M'(\nu_{\mathrm{B}}, n, \theta)} = \frac{n\gamma - n_{\mathrm{fluid}} + \rho_{\mathrm{fluid}}(\alpha + \theta(n_{\mathrm{fluid}} - n\gamma))}{\gamma^2 (n - n_{\mathrm{fluid}} + \rho_{\mathrm{fluid}}(\alpha + \theta(n_{\mathrm{fluid}} - n))} \quad (2)$$

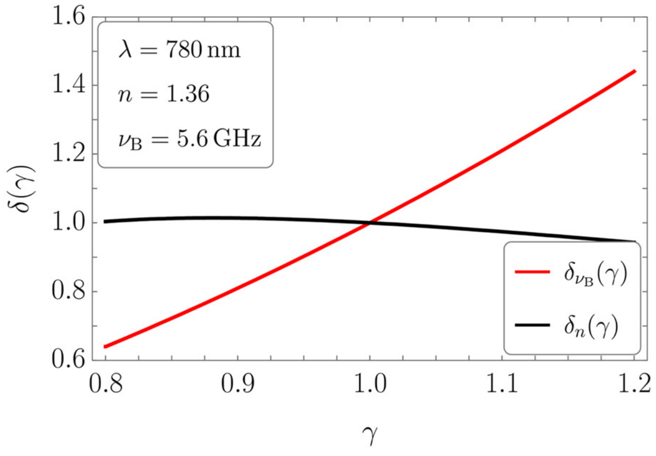

**Figure S9 | Optical diffraction tomography analysis of the zebrafish spinal lesion site, and calculations of the scaling behaviour of the longitudinal modulus and its propagated uncertainty.**

**A)** (**1**) Formula for the longitudinal modulus $M'$ after Boyd, 2008 (*2*) for Brillouin light scattering in dependence of the laser wavelength $\lambda$, the density $\rho$, the refractive index $n$, and the Brillouin frequency shift $\nu_\mathrm{B}$. (**2**) Estimation of the density for a binary mixture in dependence of the refractive index of the sample $n$, the refractive index of the inherent fluid (water) $n_\mathrm{fluid} = 1.337$, the density of the inherent fluid $\rho_\mathrm{fluid} = 0.9975\ \mathrm{g/mL}$, the refractive index increment of the dry mass content $\alpha = (0.1919 \pm 0.0030)\ \mathrm{mL/g}$, and the partial specific volume of the dry mass content $\theta = (0.743 \pm 0.010)\ \mathrm{mL/g}$ ((*3*), Table 2 'Zebrafish', values are wavelength adjusted).

**B)** Optical diffraction tomography (ODT) analysis reveals the refractive index of the lesion site in the indicated *pdgfrb*:SLRP transgenic zebrafish and their respective controls at 1 dpl. Image shown is a sagittal slice through a refractive index tomogram of the zebrafish spinal lesion site (lateral view; rostral is left). The dashed rectangle indicates the region of quantification. Each data point represents one animal. Box plots show the median, first, and third quartile. Whiskers indicate the minimum and maximum values. Scale bar: 50 µm. d, days; DOX, doxycycline; sc, spinal cord.

**C)** Analytical expressions of the Gaussian uncertainty propagation of $M'$, in $n$, $\nu_\mathrm{B}$, $\alpha$, and $\theta$ (assuming normally distributed mean values). When evaluating the ratio of uncertainty weights $\xi_n/\xi_{\nu_\mathrm{B}} \approx 0.42$, we found it to be < 1 for all control and experimental measurement values of $n$ and $\nu_\mathrm{B}$. A relative uncertainty of e.g. 10% in $n$ accounts only for a relative uncertainty in $\nu_\mathrm{B}$ of $\approx 1.1\%$.

**D)** Analytic expressions and corresponding plot for the relative changes of $M'$ in dependence of a scalar scaling parameter applied to $\nu_\mathrm{B}$ (**1**) and $n$ (**2**). For a relative scaling of e.g. 10% in any of the parameters, the relative change of $M'$ is given by $\delta_{\nu_\mathrm{B}}(\gamma = 1 \pm 0.1) \approx 1^{+0.21}_{-0.19}$ and $\delta_n(\gamma = 1 \pm 0.1) \approx 1.000^{-0.026}_{+0.014}$.

**Table S1 | Clinical data of human brain samples.**

| Case ID | Sex[1] | Age (years) | Onset (years) | Duration (years) | Side | Lobe | Diagnosis[2] | Cause[3] |
|---|---|---|---|---|---|---|---|---|
| B1 | M | 25 | 0 | 25 | left | occipital | FCD 3D | TBI |
| B2 | M | 18 | 3 | 15 | right | parietal | FCD 3D | TBI |
| B3 | M | 41 | n/a | n/a | left | occipital | FCD3D | TBI at 4 months of age |
| B4 | F | 6 | 0 | 6 | left | temp occipital | FCD 1A | 2nd surgery after 5 years |
| B5 | M | 10 | 2 | 8 | left | temp occipital | FCD 1A | 2nd surgery after 2 years |
| B6 | F | 2 | 0 | 2 | left | temp occipital | FCD 1A | 2nd surgery after 1 year |
| C1 | M | 10 | 0 | 10 | right | occipital | FCD 1A | no scar control |
| C2 | F | 18 | 13 | 5 | left | occipital | FCD 1A | no scar control |
| C3 | M | 27 | 18 | 9 | left | frontal | mMCD | no scar control |
| C4 | F | 3 | 0 | 3 | left | frontal | mMCD | no scar control |
| C5 | F | 49 | - | - | - | frontal | autopsy | aorta aneurysm |
| C6 | M | 48 | - | - | - | frontal | autopsy | heart failure |

[1] M, male; F, female; [2] FCD, focal cortical dysplasia; mMCD, mild malformation of cortical development; [3] TBI, traumatic brain injury; abbreviations: n/a, not available.

**Table S2 | Clinical and neuropathological data of human spinal cord injury cases.**

| Case ID | Age at injury (years) | Sex[1] | Injury-death interval (days) | Tissue fixation post-mortem | Type of injury[2] | Level of injury[3] | AIS grade | Diagnosis |
|---|---|---|---|---|---|---|---|---|
| BB2 | 60 | M | 111 | 5 days | C | C5 | A | Hyperextension, avulsion flakes, traumatic disc |
| BB3 | 80 | F | 17 | 6 days | C/C | C6 | B | Bilateral facet dislocation |
| BB4 | 76 | M | 34 | 13 hrs | C/C | C7 | A | Three column fracture dislocation |
| BB5 | 82 | M | 9 | 12 hrs | C | C6 | A | Bilateral facet dislocation |
| BB6 | 83 | M | 60 | 15 hrs | C/C | C4 | D | Hyperextension, avulsion flakes, traumatic disc |
| BB9 | 95 | M | 15 | 1 day | C/C | C4 | C | Hyperextension, avulsion flakes, traumatic disc |

[1] M, male; F, female; [2] C, cervical; number indicates vertebrae; [3] C/C, contusion/cyst; AIS, American Spinal Injury Association Impairment Scale.

**Table S3 | Summary of results from immunofluorescence stainings on human spinal cord samples.**

| Case ID | Increased anti-CHAD immunoreactivity in E segment[1] as compared to R/C[2] segment? | Increased anti-FMOD immunoreactivity in E segment[1] as compared to R/C[2] segment? | Increased anti-LUM immunoreactivity in E segment[1] as compared to R/C[2] segment? | Increased anti-PRELP immunoreactivity in E segment[1] as compared to R/C[2] segment? |
|---|---|---|---|---|
| BB2 | C | C | C | C |
| BB3 | R ✓ | R ✓ | R ✓ | R ✓ |
| BB4 | C | C ✓ | C ✓ | C ✓ |
| BB5 | R ✓ | R ✓ | R ✓ | R ✓ |
| BB6 | C | C | C ✓ | C |
| BB9 | R | R ✓ | R ✓ | R ✓ |

[1] E segment, epicenter-containing segment; [2] R/C segment, segment located rostral or caudal to epicenter with little to no detectable pathology, ✓ indicates an observed increased immunoreactivity in E segments as compared to C or R segments of the same case.

#### References

1. V. Tsata *et al.*, A switch in pdgfrb+ cell-derived ECM composition prevents inhibitory scarring and promotes axon regeneration in the zebrafish spinal cord. *Developmental cell* **56**, 509-524.e509 (2021).
2. R. W. Boyd, *Nonlinear Optics, Third Edition*. (Academic Press, Inc., 2008).
3. H. Zhao, P. H. Brown, P. Schuck, On the distribution of protein refractive index increments. *Biophysical journal* **100**, 2309-2317 (2011).
